# Peripheral CB1R Blockade Suppresses AKI-to-CKD Maladaptive Repair

**DOI:** 10.64898/2026.07.31.741993

**Authors:** Ariel Rothner, Liad Hinden, Aviram Kogot-Levin, Shridhar Betkar, Elisheva Benkovitz, Amani Zoabi, Anna Permyakova, Asaf Kleiner, Vladislav Nesterenko, Alina Nemirovski, Ifat Abramovich, Bella Agranovich, Inbar Plaschkes, Eyal Gottlieb, Katherine Margulis, Gil Leibowitz, Joseph Tam

## Abstract

**Background:** Acute kidney injury (AKI) frequently progresses to chronic kidney disease (CKD), yet mechanisms governing this transition remain poorly understood. The endocannabinoid system (ECS), particularly cannabinoid-1 receptor (CB1R), regulates inflammation and metabolism in various organs, but its role in post-AKI maladaptive repair is less established.

**Methods:** We analyzed CB1R expression in kidney biopsies from pre- and post-transplant recipients and in murine AKI models (ischemia-reperfusion injury [IRI] and folic acid [FA]-induced AKI). Peripheral CB1R blockade was evaluated in FA-AKI model and in human primary kidney proximal tubule cells (hKPTCs). Spatial metabolomics, semi-targeted metabolomic profiling, and gene and protein expression characterized molecular mechanisms.

**Results:** CB1R expression was increased in kidneys undergoing maladaptive repair in both humans and mice, but remained unchanged during acute injury. In the FA-induced AKI model, the ECS showed stage-specific alterations, with temporal and spatial fluctuations in endocannabinoid levels and their enzymatic regulators. Peripheral CB1R blockade during the repair phase preserved kidney function, reduced injury, and maintained systemic glucose homeostasis. Metabolomic and molecular analyses revealed that CB1R blockade restored dysregulated arginine metabolism and reduced AKT/NF-κB-p65 pathway in post-AKI kidneys, linking CB1R activation to inflammatory signaling. In hKPTCs, 2-AG-induced activation of CB1R increased VCAM1 expression, a failed-repair marker, while its antagonism reduced TNFα/2-AG-induced expression of pro-inflammatory adhesion molecules, chemokines, cytokines, and arginine metabolism enzymes.

**Conclusions:** CB1R overactivation drives AKI-to-CKD progression by promoting inflammatory signaling and metabolic dysregulation. Peripheral CB1R blockade during the repair phase represents a novel therapeutic strategy to prevent maladaptive repair and CKD development after AKI. These findings establish CB1R as a phase-specific therapeutic target for post-AKI intervention.

**Translational Statement:** Peripheral CB1R antagonists offer a first-in-class therapeutic strategy to halt progression from acute kidney injury (AKI) to chronic kidney disease (CKD) by selectively targeting maladaptive tubular repair. By blocking CB1R signaling specifically in the kidney, these agents attenuate inflammation, metabolic dysregulation, and fibrogenic pathways that drive failed repair, while sparing central nervous system CB1R and thereby minimizing neuropsychiatric adverse effects. This phase-specific, peripherally restricted approach supports the development of peripheral CB1R antagonists as a viable translational therapy to improve long-term renal outcomes after AKI.

## Introduction

Acute kidney injury (AKI) is a common and serious clinical syndrome affecting millions worldwide and representing a major cause of morbidity and mortality^1^. Studies over recent decades demonstrate that AKI survivors face substantially elevated risk of chronic kidney disease (CKD) and end-stage kidney disease^2–4^. The transition from AKI to CKD is driven by maladaptive repair, characterized by tubular epithelial loss, extracellular matrix accumulation, and persistent inflammation, culminating in renal fibrosis and functional decline^5,6^. Renal inflammation, particularly sustained activation of immune cells and pro-inflammatory transcription factors such as nuclear factor-κB (NF-κB), plays a central role in driving maladaptive repair and CKD progression^7,8^.

The endocannabinoid system (ECS), consisting of G-protein-coupled cannabinoid receptors (CB1R and CB2R), their endogenous ligands (endocannabinoids/eCBs), and metabolic enzymes, plays a key role in kidney health and disease^9^. ECS overactivation has been implicated in chronic kidney diseases^10–12^, with CB1R blockade shown to alleviate diabetic nephropathy, obesity-related CKD, and unilateral urethral obstruction (UUO) in preclinical models^13–19^. However, CB1R’s role in AKI appears context- and phase-dependent. Studies examining CB1R during acute injury report conflicting results, both protective and deleterious effects depending on injury model^20–23^, and have primarily focused on the acute phase. Yet, the role of CB1R in post-AKI maladaptive repair, a key factor shaping long-term outcomes, remains largely unexplored.

Here, we hypothesized that CB1R plays a phase-specific role in AKI progression, favoring maladaptive repair over acute injury. We characterized ECS dynamics across disease stages in human kidney transplant recipients and two complementary murine AKI models (ischemia-reperfusion injury [IRI] and folic acid [FA]-induced AKI). CB1R expression increased selectively in failed-repair proximal tubular cells (FR-PTCs) during late-stage injury, while remaining unchanged during acute injury. Using peripheral CB1R blockade in *in-vitro* and *in-vivo* settings, we demonstrate that CB1R drives post-AKI maladaptive repair by activating inflammatory (NF-κB) and metabolic (arginine dysregulation) pathways. These findings identify CB1R as a therapeutic target to prevent AKI-to-CKD progression and establish the rationale for phase-specific intervention strategies.

## Methods

**Supplemental Information** includes detailed descriptions of antibodies, primers, reagents, and extended protocols.

### Transcriptomic analyses

#### Clinical Dataset

Bulk RNA-seq data from 42 kidney transplant recipients (KTRs; GSE162805) were analyzed at four time points: pre-implantation, post-reperfusion, 3 months, and 12 months post-transplant^24^. As described in *Cippa et. al*, samples at 3 and 12 months were clustered using the Walktrap algorithm into communities reflecting injury/fibrosis (Community A) or healthier profiles (Community B), correlating with kidney function and fibrosis scores. Differential expression was assessed using DESeq2 (FDR < 0.1). *Mouse Datasets:* Bulk RNA-seq data from bilateral ischemia-reperfusion injury (IRI; GSE98622) were analyzed across multiple time points (2 hours to 12 months post-injury) with sham-operated controls. Single-nucleus Assay for Transposase-Accessible Chromatin using sequencing (snATAC-seq)^7^ and single-nucleus RNA sequencing (snRNA-seq)^25^ from post-IRI kidneys were accessed via the Kidney Interactive Transcriptomics database (https://humphreyslab.com/SingleCell/).

### Murine AKI studies

All procedures were approved by the Hebrew University of Jerusalem Institutional Animal Care and Use Committee (Ethics approval MD-19-15935, MD-22-17030) and adhered to ARRIVE guidelines. For the FA-induced AKI model, male C57BL/6J mice received an i.p. injection of 240 mg/kg FA or vehicle (0.3 M sodium bicarbonate, injection volume of 10 µL/g, an average of 270 µL per mouse). Urine was collected at 2, 14, and 28 days using the CCS2000 Chiller System, and mice were euthanized under anesthesia for tissue collection. For CB1R blockade studies, mice received daily oral gavage of JD5037 (3 mg/kg) or placebo (1% Tween80, 4% DMSO, 95% saline) for 2- or 14-day post-FA. A healthy cohort received JD5037 (3 mg/kg, i.p.) or placebo for 7 days. For the unilateral IRI with delayed nephrectomy (uIRI), male C57Bl/6J mice underwent renal pedicle clamping for 21 min to induce IRI to the left kidney on day 0, followed by a right kidney nephrectomy on day 14. For bilateral IRI (bIRI), male C57Bl/6J mice underwent renal pedicle clamping for 28 min on day 0 in both kidneys. Sham control mice underwent identical surgical procedures, with the exception of ischemic clamping. On days 15 for uIRI and 14 for bIRI, mice were euthanized by cervical dislocation under anesthesia; trunk blood and kidneys were collected, and samples were snap-frozen for further analysis.

### Biochemical analyses

Serum BUN, creatinine, glucose, and urine parameters were analyzed using the Cobas C-111 analyzer (Roche). Serum NGAL was measured by ELISA (Abcam). Creatinine clearance was calculated as: (Urine creatinine × Urine volume)/(Serum creatinine × 24 h]. FAAH activity in kidney lysates was assessed using the FAAH Activity Assay Kit (Abcam).

### Metabolomics

LC-MS-based metabolomic profiling was performed on kidney, liver, and serum samples. Samples were processed using a Dionex Ultimate 3000 UPLC coupled to an Orbitrap Q-Exactive Mass Spectrometer (Thermo Fisher) with a ZIC-pHILIC column and a 15-min gradient. Metabolites were identified using a validated in-house MS library and analyzed with MetaboAnalyst 5.0.

### Endocannabinoid quantification

Endocannabinoids (AEA, 2-AG) and arachidonic acid (AA) were extracted from kidney and urine using stable isotope dilution LC-MS/MS on a QTRAP® 6500+ (Sciex). Samples were homogenized in Tris buffer, extracted with chloroform/methanol, and analyzed by multiple reaction monitoring (MRM).

### Spatial metabolomics (DESI-MSI)

Kidney sections (20 µm) were analyzed using a Xevo G2-XS Q-TOF mass spectrometer (Waters) at 50 µm spatial resolution. Data were normalized by ion intensity within the selected ROI.

### Histopathology and Immunohistochemistry

Paraffin-embedded kidney sections (4 µm) were either stained with PAS and Masson’s trichrome (Abcam) to assess tubular injury and fibrosis or stained for F4/80 to assess macrophage infiltration. Quantification was performed using Image Pro Plus 6.0.

### Gene expression

RNA was extracted from the kidney cortex, and qPCR was performed using SYBR Green Supermix (Bio-Rad). Expression was normalized to *S16* or *Gapdh*. Primer sequences are provided in **Supplementary Table 3**.

### Protein expression

Kidney lysates were processed in RIPA buffer, resolved via SDS-PAGE, and transferred to PVDF membranes. Proteins were detected with primary antibodies (listed in **Supplementary Table 4**) and quantified by densitometry using Bio-Rad CFX Manager software, normalized to total protein or housekeeping protein controls.

### Cell culture

Primary human kidney proximal tubule cells (hKPTCs; Lonza; Cat #CC-2253) were cultured in REGM BulletKit medium (Lonza; Cat #CC-3191 & CC-4127). After an overnight starvation in serum-free medium, hKPTCs were pre-incubated with/without JD5037 (100 nM) for one hour and supplemented with different treatments, including TNFα, noladine ether, or their combination for an additional 6 hours. The cells were then collected for RNA extraction and qPCR.

### Statistical analysis

GraphPad Prism v10 was used for statistical analyses. Comparisons between two groups used Student’s t-test or Mann-Whitney U-test. Multiple group comparisons used a one-way ANOVA with Tukey’s or Dunnett’s post hoc tests. Pearson correlation and logistic regression were applied for predictive modeling. Significance was set at p < 0.05.

## Results

### CB1R is selectively upregulated during renal maladaptive repair

Ischemia-reperfusion is an inevitable consequence of kidney transplantation and can contribute to progressive allograft dysfunction by promoting maladaptive repair, chronic inflammation, and fibrosis. To investigate ECS involvement in post-transplant kidney injury, we retrospectively analyzed published bulk RNA-sequencing data from KTRs at four time points: pre-implantation, post-reperfusion, 3 months, and 12 months post-transplant^24^ (**Figure 1a**). Samples collected immediately before (Pre) and after (Post) renal reperfusion displayed no differences in gene expression of the cannabinoid receptors (**Supplementary Figure 1a, b**) or the eCB metabolic enzymes (**Supplementary Figure 1c-g**). At 3- and 12-month post-transplantation, patient samples clustered into two distinct communities: Community A, characterized by a maladaptive repair transcriptional profile, reduced renal function, and increased fibrosis; and Community B, displaying a healthier recovery profile (**Figure 1a**).

**Figure 1.**
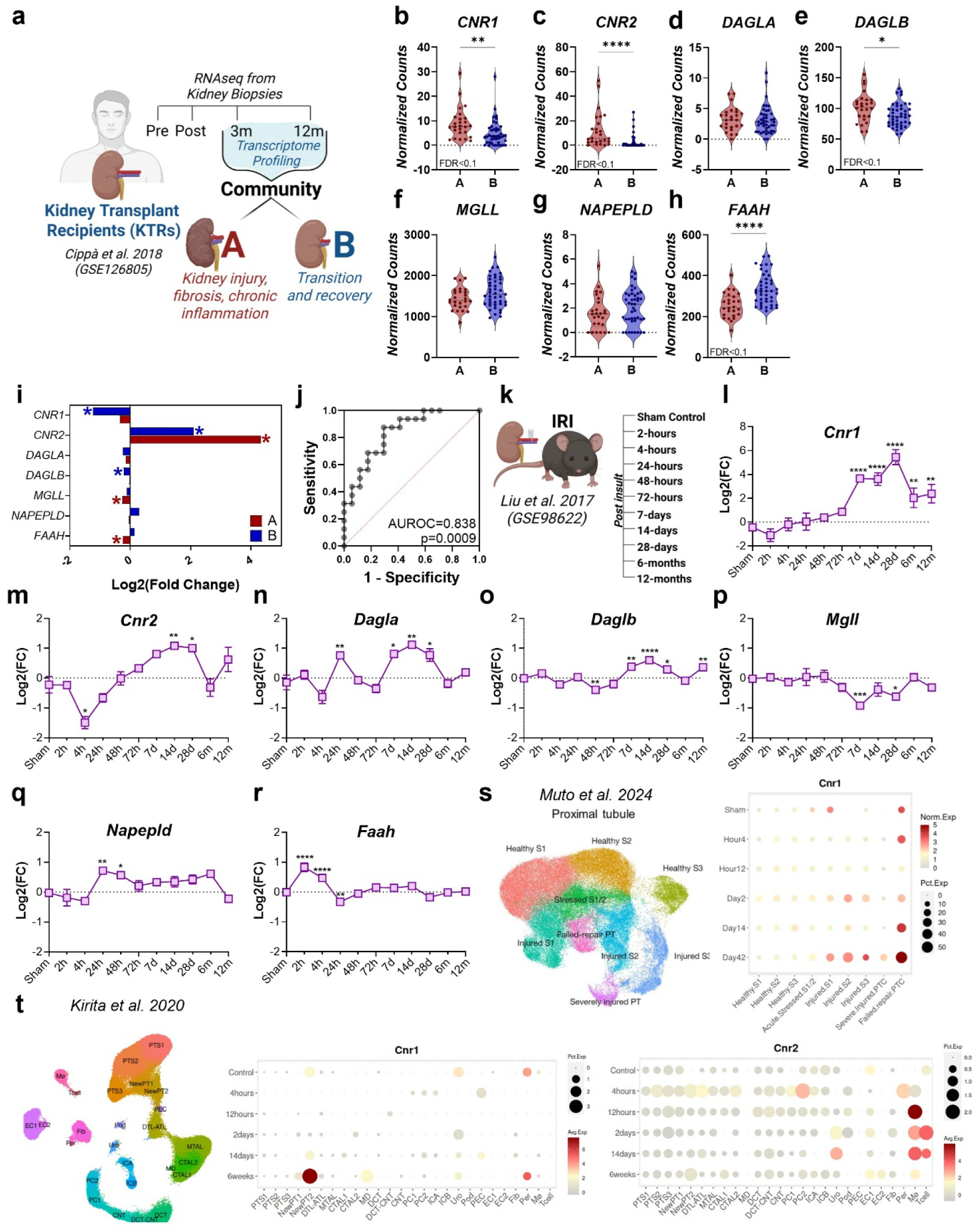
Enhanced CB1R expression and activation in post-AKI maladaptive repair. **(a)** RNA-Seq of mRNA extracted from whole kidneys of 42 KTR at 3- and 12-month post-reperfusion (GSE126805). At follow-up, patients were stratified into two communities based on transcription profiles: Community A (red), showing injury, inflammation, and fibrosis, and Community B (blue), representing healthy recovery, as previously described^24^. **(b-c)** Gene expression levels of cannabinoid receptors *CNR1* **(b)** and *CNR2* **(c)** in these two groups; **(d-h)** Expression of key eCB metabolic enzymes: *DAGLA* **(d)**, *DAGLB* **(e)**, *MGLL* **(f)**, *NAPEPLD* **(g)**, and *FAAH* **(h)**. Comparison between groups was performed by Mann-Whitney test (*p < 0.05, **p < 0.01, ***p < 0.001, ****p < 0.0001); differentially expressed genes by DESeq2 (FDR < 0.1) are indicated. **(i)** Changes in gene expression at 3- and 12-month post-transplant compared to immediate post-reperfusion samples, shown by community; *FDR < 0.1. **(j)** ROC curve (AUROC) of a logistic regression for *CNR1* expression predicting kidney fibrosis (ci > 0 vs. ci = 0) at 12 months post-transplantation. **(k)** RNA-sequencing of whole mouse kidney at multiple time points after bilateral IRI (n = 3-4/group) compared to sham control (analyzed at 4-hour, 24-hour, 12-month; n=9) from NCBI GEO accession GSE98622 showing **(l-r)** expression profiles for ECS-related genes, data represent the mean ± SEM. Comparisons by one-way ANOVA with Dunnett’s post-hoc test vs sham (*p < 0.05, **p < 0.01, ***p < 0.001, ****p < 0.0001). Mouse kidneys following bilateral IRI were collected at various time points as described in Kirita et al., 2020^25^: **(s)** snATAC-seq subclustering of mouse PTCs by cell subtype and chromatin accessibility at the *Cnr1* locus. **(t)** snRNA-seq UMAP plots of *Cnr1* and *Cnr2* transcript levels across kidney cell clusters, with New-PT1 and New-PT2 denoting injured PTCs. *Abbreviations:* KTR, kidney transplant recipients; *CNR1/2*, cannabinoid-1/-2 receptor; *DAGLA/B*, diacylglycerol lipase A/B; *MGLL*, monoacylglycerol lipase; *NAPEPLD*, *N*-acyl phosphatidylethanolamine phospholipase D; *FAAH*, fatty acid amide hydrolase; ECS, endocannabinoid system; IRI, ischemia reperfusion injury; ATL, thin ascending limb of loop of Henle; Bil, bilateral; CNT, connecting tubule; CPC, principle cells of collecting duct in cortex; CTAL, thick ascending limb of loop of Henle in cortex; DCT, distal convoluted tubule; DTL, descending limb of loop of Henle; EC, endothelial cells; Fib, fibroblasts; ICA, type A intercalated cells of collecting duct; ICB, type B intercalated cells of collecting duct; MD, macula densa; Mø, macrophages; MPC, principle cells of collecting duct in medulla; MTAL, thick ascending limb of loop of Henle in medulla; PEC, parietal epithelial cells; Per, pericytes; Pod, podocytes; PT-S1, S1 segment of proximal tubule; PT-S2, S2 segment of proximal tubule; PT-S3, S3 segment of proximal tubule; Uro, urothelium.

Community A showed significantly higher expression of cannabinoid receptors *CNR1* and *CNR2* (**Figure 1b, c**) and evidence of enhanced endogenous ligand 2-arachidonoyl glycerol (2-AG) signaling. This was indicated by increased expression of *DAGL* (the 2-AG-synthesizing enzyme diacylglycerol lipase) and unchanged expression of MGLL (the 2-AG-catabolic enzyme monoacylglycerol lipase), collectively suggesting elevated 2-AG production (**Figure 1d-f**). Similarly, enhanced abundance of the canonical eCB anandamide (AEA) was indicated in Community A, supported by consistent expression of its synthesizing enzyme *N*-acyl phosphatidylethanolamine phospholipase D (*NAPEPLD*) and reduced expression of its catabolic enzyme fatty acid amide hydrolase (*FAAH*) (**Figure 1g, h)**. Importantly, temporal analysis revealed distinct ECS tone dynamics between groups: Community B (healthy recovery) showed downregulation of *CNR1* and *DAGLB*, suggesting a decline in ECS signaling, whereas Community A exhibited decreased catabolic enzymes (*MGLL* and *FAAH*), suggesting sustained or elevated ECS tone (**Figure 1i**). Critically, *CNR1* expression was associated with the severity of renal fibrosis. Logistic regression and receiver operating characteristic (ROC) analysis demonstrated *CNR1* expression at 12 months significantly discriminates the presence of interstitial fibrosis (ci > 0; **Figure 1j**), supporting previous findings where CB1R expression correlated with chronic allograft dysfunction and renal fibrosis^26^.

We next evaluated bulk RNA-seq data from a murine bIRI model at multiple time-points post-injury^27^ (2 hours to 12 months; **Figure 1k**). Paralleling the KTR data, both cannabinoid receptor expression increased only during later stages post-IRI and persisted long-term, with *Cnr1* levels rising over 30-fold between 7 days and 12 months post-injury (**Figure 1l, m**). Additionally, enzymes involved in 2-AG metabolism showed significant alterations during the maladaptive repair phase, with elevated *Dagl* expression and suppressed *Mgll*, indicating enhanced 2-AG production (**Figure 1n-p**). In contrast, AEA metabolic enzymes changed within the first 48 hours post-injury (**Figure 1q, r**), supporting enhanced AEA turnover during the initial insult and distinguishing acute injury signaling from repair-phase eCB responses.

Single-nucleus transcriptomic and epigenomic analyses across studies showed that injury induces a subset of vascular cell adhesion molecule 1 (*Vcam1*)-expressing proximal tubule cells (PTCs) with distinct proinflammatory, profibrotic, and epigenetically altered signatures. This subset, termed failed-repair PTCs (FR-PTCs), persists post-injury and is suggested to promote CKD progression^7,25,28^. Analysis of post-IRI kidneys by single-nucleus ATAC-seq showed that *Cnr1* exhibits late-stage increases in chromatin accessibility specifically within FR-PTCs (**Figure 1s**), establishing epigenetic priming for CB1R expression in this pathogenic subset. Single-nucleus RNA-seq confirmed that *Cnr1* expression is selectively enriched within NewPT1 and NewPT2 (denoting FR-PTCs) (**Figure 1t**), positioning CB1R as a potential driver of maladaptive repair^7,25,28^. Meanwhile, *Cnr2* expression was enriched mainly in immune cells (**Figure 1t**). Together, the temporal dissociation, unchanged during acute injury but progressively upregulated during maladaptive repair, demonstrates a phase-specific pathogenic role, positioning CB1R as a therapeutic target for preventing AKI-to-CKD progression.

### Peripheral CB1R blockade during the repair phase mitigates renal damage

Given CB1R’s selective upregulation during maladaptive repair, we next tested whether peripheral CB1R blockade could prevent disease progression. To enable longitudinal pharmacological intervention, we employed a folic acid (FA)-induced AKI model that recapitulates the AKI-to-CKD transition, characterized by transient renal dysfunction yet persistent tubular injury, inflammation, and fibrosis (**Supplementary Figure 2**).

Mice received the peripherally restricted CB1R inverse agonist JD5037 (3 mg/kg, p.o) or vehicle for 2- or 14-days following FA-AKI induction (**Figure 2a**). Consistent with CB1R’s lack of upregulation during acute injury, JD5037 did not attenuate AKI severity at day 2 (**Supplementary Figure 3**). However, by day 14, when CB1R expression peaks in this model (**Figure 3a, c, d**), CB1R blockade significantly reduced BUN, serum creatinine, and renal expression of clusterin (*Clu*) and neutrophil gelatinase-associated lipocalin (NGAL/*Lcn2*) (**Figure 2b-f**) compared to vehicle-treated controls. Histological analysis revealed that peripheral CB1R blockade preserved tubular architecture and reduced injury severity by ameliorating tubular necrosis, dilation, and brush border loss, with less effect on cast formation (**Figure 2g-k**).

**Figure 2.**
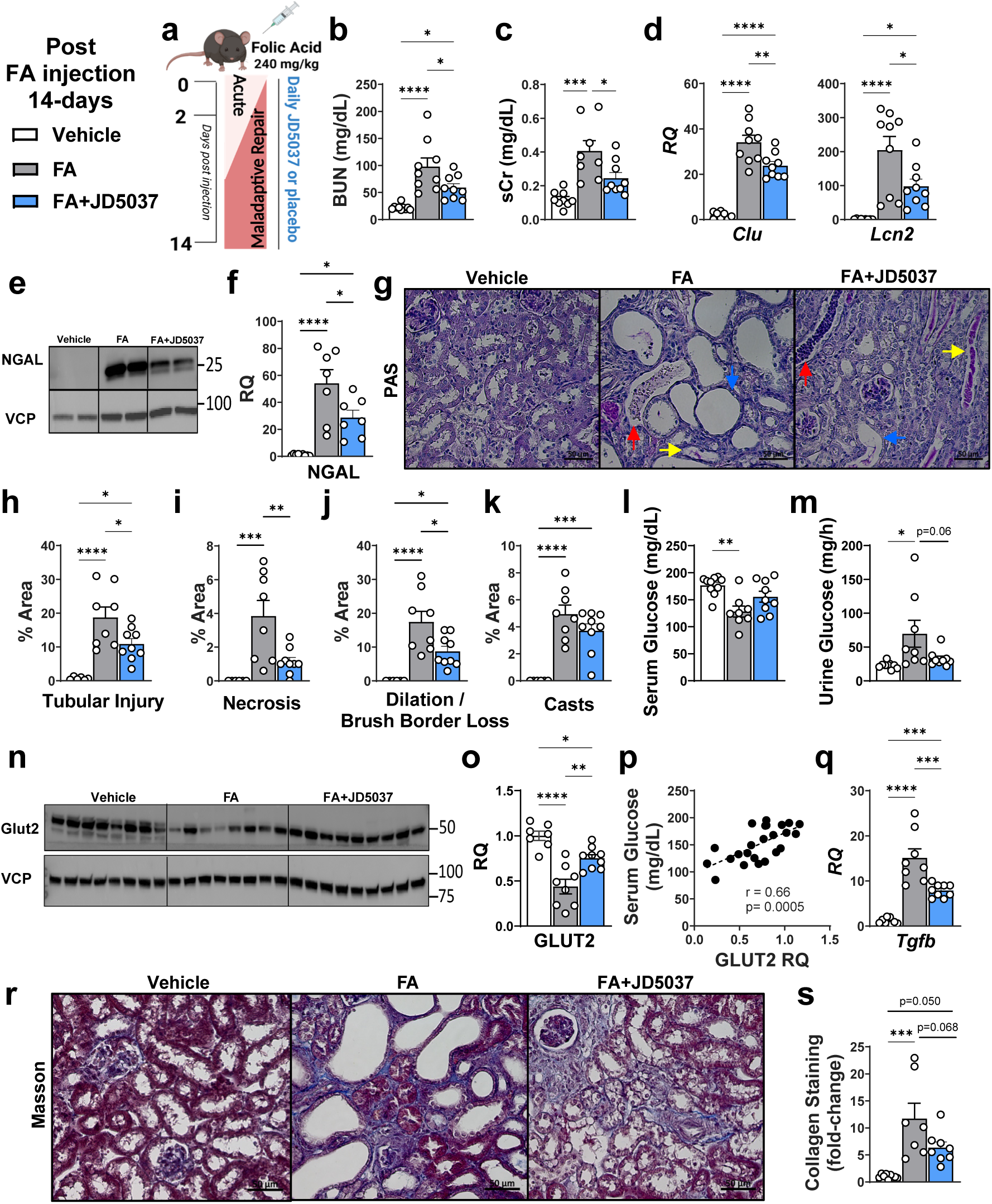
Peripheral CB1R inverse agonism ameliorates post-AKI maladaptive repair in the FA model. Daily oral treatment with the peripherally restricted CB1R inverse agonist JD5037 (3 mg/kg) or placebo was orally administered for two weeks following FA-induced AKI (240 mg/kg) or vehicle (0.3 M sodium bicarbonate) in mice (n = 9-10 per group). **(b)-(c)** Kidney function markers: serum BUN and creatinine (sCr) following treatment. **(d)-(f)** Renal injury markers: transcript expression (*Clu* and *Lcn2*) and immunoblotting/quantification for NGAL, normalized to VCP. **(g)** Representative kidney histopathological assessment (PAS stains) showing tubular necrosis (red arrow), cast formation (yellow arrow) and tubular dilation/brush border loss (blue arrow). Scale bar: 50 μm, magnification 40×. **(h-k)** Quantitative analysis of total kidney injury, and breakdown of necrosis, brush border loss, and cast area. **(l-m)** serum and urinary glucose levels. **(n-o)** GLUT2 protein expression by immunoblot and quantification, normalized to VCP (n = 7-9 per group). **(p)** Linear regression analysis of serum glucose vs. GLUT2 expression. **(q)** Kidney *Tgfb* transcript expression. **(r-s)** Masson’s trichrome staining and quantification of collagen deposition. Scale bar: 50 μm, magnification 40×. Data represent the mean ± SEM (n = 7-10 per group). Differences among experimental groups were analyzed using one-way ANOVA followed by Tukey’s post-hoc test. *p < 0.05, **p < 0.01, ***p < 0.001, ****p < 0.0001. *Abbreviations:* AKI, acute kidney injury; CB1R, cannabinoid-1 receptor; FA, folic acid; BUN, blood urea nitrogen; sCR, serum creatinine; *Clu*, clusterin; *Lcn2*/NGAL, neutrophil gelatinase-associated lipocalin; VCP, valosin-containing protein; GLUT2, glucose transporter 2; *Tgfb*, transforming growth factor beta.

**Figure 3.**
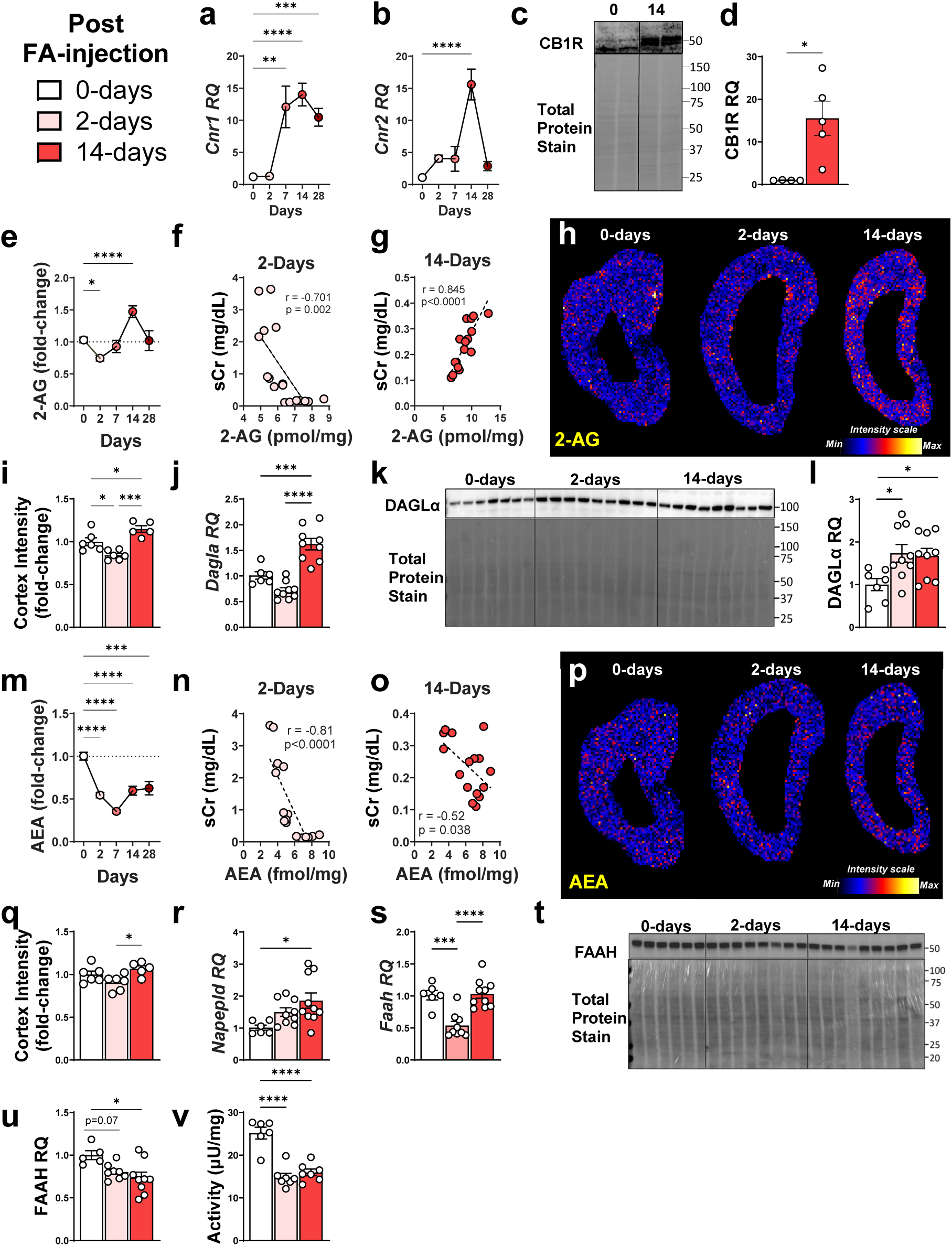
Distinct spatial and temporal ECS response following FA-induced AKI. Male C57Bl/6J mice were injected i.p. with FA (240 mg/kg) to induce AKI and assessed at baseline (0 days), and at 2-, 7-, 14-, and 28-day post-injection (n = 6-11 mice per group). **(a, b)** Renal transcript expression of *Cnr1* and *Cnr2* over time. **(c, d)** Renal CB1R protein by Western blot (normalized to total protein) and its quantification comparing 0- and 14-day post-FA. **(e)** Abundance of 2-AG in the renal cortex relative to baseline. **(f, g)** Linear regression and Pearson correlation between sCr and 2-AG levels at 2 and 14 days. **(h, i)** DESI-MSI mapping 2-AG in kidneys at different time points, with ROI in cortex, presented as heatmaps alongside quantification (n = 5-6 per group). **(j-l)** Renal *Dagla*/DAGLα transcript expression, protein by Western blot, and quantification (normalized to total protein). **(m)** Abundance of AEA in renal cortex relative to baseline. **(n, o)** Linear regression and Pearson correlation between sCr and AEA at 2- and 14-day post-FA. **(p, q)** DESI-MSI mapping of AEA and quantification in kidneys at different time points, with ROI in cortex, presented as heatmaps alongside quantification (n = 5-6 per group). **(r, s)** Renal transcript expression of *Napepld* and *Faah*. **(t, u)** FAAH protein expression and quantification (normalized to total protein). **(v)** FAAH enzymatic activity. Data represent the mean ± SEM. For statistical analysis: Student’s t-test (two groups), one-way ANOVA with Tukey’s post-hoc test (three groups), or Dunnett’s test (more than three groups) were used as appropriate. *p < 0.05, **p < 0.01, ***p < 0.001, ****p < 0.0001. *Abbreviations:* ECS, endocannabinoid system; AKI, acute kidney injury; FA, folic acid; *Cnr1*/CB1R, cannabinoid-1 receptor; *Cnr2*, cannabinoid-2 receptor; VCP, valosin-containing protein; 2-AG, 2-arachidnoylglycerol; sCR, serum creatinine; DESI-MSI, desorption ionization mass spectrometry imaging; ROI, region of interest; *Dagla*/DAGLα, diacylglycerol lipase; *Mgll*/MAGL, monoacylglycerol lipase; AEA, anandamide; *Napepld*, *N*-acyl phosphatidylethanolamine phospholipase D; *Faah*/FAAH, fatty acid amide hydrolase.

Peripheral CB1R blockade also preserved glucose homeostasis, maintaining serum glucose levels and preventing glucosuria (**Figure 2l, m**), and restored glucose transporter-2 (GLUT2) expression (**Figure 2n, o**), which correlated with serum glucose levels (r = 0.66; p = 0.0005; **Figure 2p**). These metabolic effects were independent of sodium glucose transporter-2 (SGLT2) expression or correction of the metabolic disturbances in gluconeogenesis/glycolysis metabolites or enzyme expression (**Supplementary Figure 4a-g**), identifying GLUT2 as a key mediator of CB1R antagonism’s metabolic benefits. Notably, disruption of these transporters is a feature of clinical maladaptive repair, as their expression was significantly lower in KTRs from Community A than in their healthy counterparts (Community B; **Supplementary Figure 1h-i**).

Treatment with JD5037 also reduced profibrotic transforming growth factor (*Tgfb*) expression and reduced collagen deposition (**Figure 2q-s**), though anti-fibrotic effects may be more pronounced with extended treatment duration, given that fibrosis peaks at day 28 in this model (**Supplementary Figure 2g, k**). Overall, these findings demonstrate that peripheral CB1R blockade during the repair phase, when CB1R is upregulated, prevents functional decline and preserves tubular integrity, supporting phase-specific therapeutic intervention.

### Endocannabinoid system dynamics during AKI and maladaptive repair

To contextualize the therapeutic effects of CB1R blockade, we next examined ECS dynamics across stages of AKI and maladaptive repair using multiple *in-vivo* renal injury models. In FA-induced AKI, gene expression of cannabinoid receptors remained unchanged in the acute phase, but gradually increased during maladaptive repair, peaking at 14-days (**Figure 3a, b**). This elevation in CB1R expression at 14-days was confirmed at the protein level using a validated antibody (**Figure 3c, d**). This Cayman CB1R antibody was validated using brain tissue lysates from wild-type and CB1R knockout mice (**Supplementary Figure 5a**). Enhanced gene expression of the cannabinoid receptors, specifically in the later stages, was also demonstrated in uIRI and bIRI murine models (**Supplementary Figure 6a-c, q-s**).

Next, we examined eCB levels in the different injury stages. 2-AG, the predominant renal eCB, exhibited stage-specific changes. In FA-AKI renal cortical 2-AG levels initially decreased at day 2 but increased significantly to 50% above pre-injury levels by day 14 (**Figure 3e**, **Supplementary Table 1**). Importantly, 2-AG levels correlated with kidney dysfunction at both 2 and 14 days; however, the direction of this relationship reversed between the 2 time points, shifting from negative at day 2 to positive at day 14 (**Figure 3f, g**). Spatial metabolomics by DESI-MSI revealed a dynamic distribution of 2-AG, confirming the cortical reduction at day 2 and increased levels at day 14 (**Figure 3h-i**). We also observed medullary 2-AG accumulation during acute injury (day 2) that became evenly distributed by day 14 (**Supplementary Figure 5b-d**). This was accompanied by decreased urinary 2-AG levels, with a similar trend for its degradation product, AA (**Supplementary Figure 5e-f**). This reduced urinary 2-AG was observed despite increased urinary output in injured mice and was negatively correlated with serum creatinine levels (**Supplementary Figures 2e, 5g**). These 2-AG alterations were driven by increased cortical expression of its synthesizing enzyme *Dagla*/DAGLα at day 14 (**Figure 2j-l**), while its catabolic enzyme *Mgll*/MAGL remained unchanged (**Supplementary Figure 5h-j**), potentially accounting for spatial 2-AG dynamics. Similar temporal changes were also observed in the IRI-AKI models, with elevated 2-AG abundance at 15-days after uIRI (**Supplementary Figure 6d**) and corresponding alterations in anabolic and catabolic enzyme expression in both uIRI and bIRI models (**Supplementary Figure 6e-g, t-v**).

Similar to 2-AG in the FA-AKI model, AEA levels decreased acutely in the kidney cortex, remained low through day 14, and correlated inversely with serum creatinine at both time points (**Figure 3m-o**, **Supplementary Table 1**). While spatial analysis did not reveal reduced AEA levels at 2 days, cortical AEA was higher at day 14 compared to day 2 post-injury (**Figure 3p-q**). The injury also altered renal AEA spatial organization, shifting from uniform distribution in healthy kidneys to medullary accumulation at day 2 post-injection (**Supplementary Figure 5k-m**). These alterations in AEA levels occurred despite compensatory upregulation of its synthesizing enzyme *Napepld* (**Figure 3r**) and decreased *Faah*/FAAH expression and activity (**Figure 3s-v**), suggesting involvement of alternative AEA metabolic pathways^13^ or enhanced substrate depletion. Likewise, renal AEA was depleted at both acute and chronic stages following uIRI (**Supplementary Figure 6h**), and varying alterations in its metabolic enzyme expression (**Supplementary Figure 6i-j, w-x**). Altogether, these stage-specific ECS alterations support phase-specific targeting of CB1R, with the late-stage elevation of 2-AG reinforcing CB1R overactivation as a driver of post-AKI maladaptive repair.

### Peripheral CB1R blockade normalizes disrupted arginine metabolism during maladaptive repair

To elucidate the molecular mechanisms underlying the protective effects of peripheral CB1R blockade in maladaptive repair, we performed metabolomic profiling of kidneys from FA-AKI mice treated with JD5037 or vehicle. Post-AKI kidneys displayed profound metabolic alterations, with over half of the detected metabolites differing from those of healthy controls (**Figure 4a**). Principal component analysis showed that the metabolomic profiles clustered distinctly with JD5037, shifting the post-AKI signature toward a healthy phenotype (**Figure 4b**) via modulating key metabolites important to the arginine biosynthesis and metabolism pathway (**Figure 4c**).

**Figure 4.**
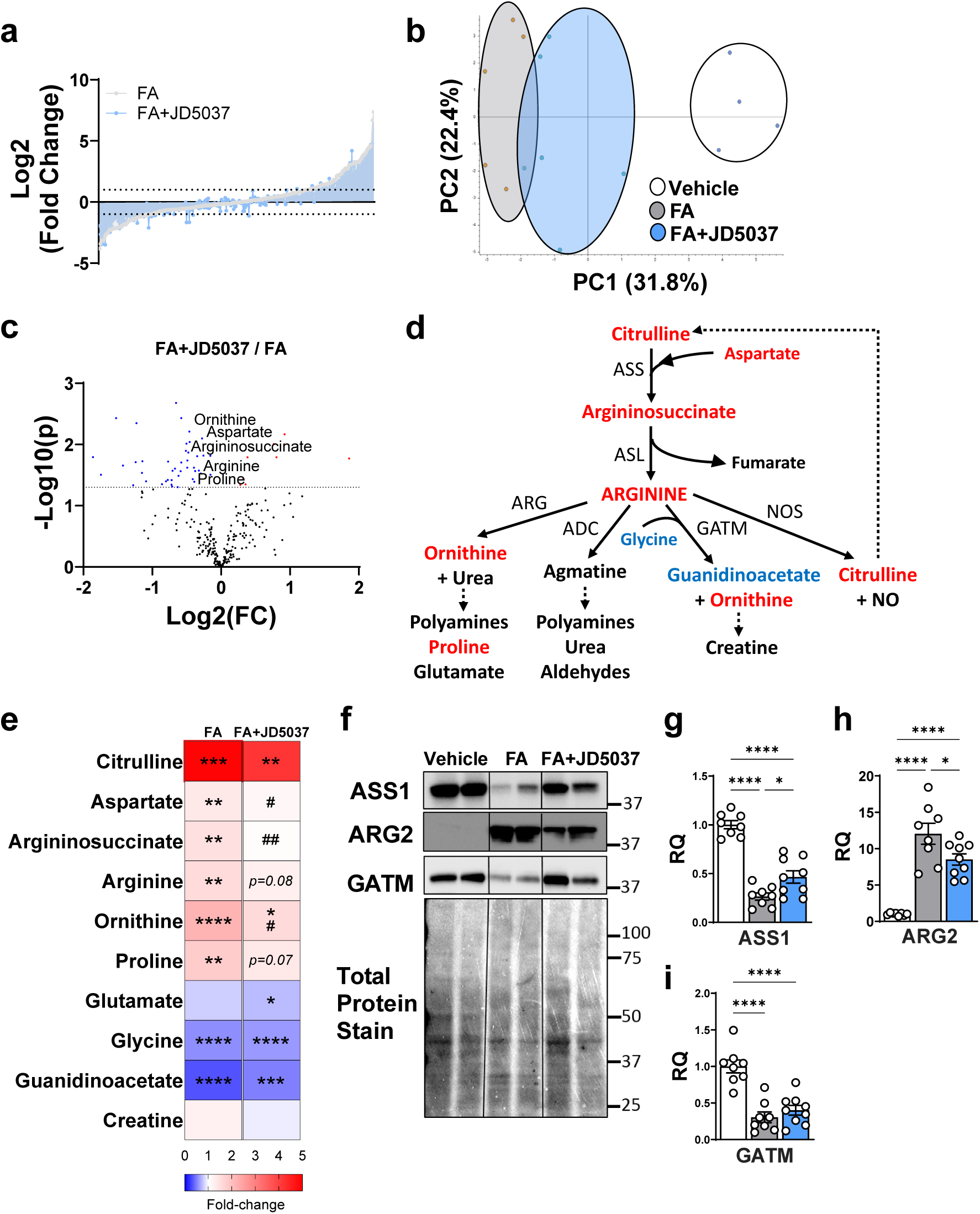
Peripheral CB1R blockade post-AKI induces a distinct metabolomic profile and reduces dysregulation of the arginine pathway. LC-MS/MS-based metabolomic profiling was performed on kidneys collected two weeks after FA-induced AKI (240 mg/kg) in mice orally treated daily with the peripherally restricted CB1R inverse agonist JD5037 (3 mg/kg) or vehicle (n = 4-6 per group). **(a)** Relative change in 300 semi-targeted detected metabolites, comparing FA and FA+JD5037 groups to vehicle control. **(b)** PCA of the total untargeted metabolome. **(c)** Volcano plot depicting metabolite changes between the FA+JD5037 and non-drug-treated FA groups; metabolites significantly increased (red) or decreased (blue) in the drug group are shown. **(d)** Schematic representation of the arginine biosynthesis pathway, highlighting metabolites significantly increased (red) or decreased (blue) after FA-AKI. **(e)** Heatmap showing relative levels of metabolites in the arginine biosynthesis pathway for all groups (*indicates comparison to vehicle; #indicates comparison to FA group; ^#^p<0.05, ^##^p < 0.01, or as specified). **(f)** Immunoblot analysis of renal arginine metabolic enzymes and **(g-i)** quantification of protein expression (n = 8-9 per group). Data represent the mean ± SEM. Statistical analysis employed one-way ANOVA with Tukey’s post-hoc test. *p < 0.05, **p < 0.01, ***p < 0.001, ****p < 0.0001. *Abbreviations:* CB1R, cannabinoid-1 receptor; FA, folic acid; AKI, acute kidney injury; ASS, arginosuccinate synthase; ASL, arginosuccinate lyase; ARG, arginase; ADC, arginine decarboxylase; GATM, L-Arginine:glycine amidinotransferase; NOS, nitric oxide synthase; NO, nitric oxide.

Arginine and its derivatives play crucial roles in renal inflammation, tissue repair, and fibrogenesis, processes contributing to maladaptive repair^29^. The kidney is a major site of arginine biosynthesis and metabolism, influencing various metabolic pathways that depend on specific arginine-catabolic enzymes^30,31^ (**Figure 4d**). In fact, pathway enrichment analysis highlighted disruptions in these pathways during both the acute (2-day) and maladaptive repair (14-day) stages, with noticeably different patterns among metabolites (**Supplementary Figure 7, Supplementary Data File**). Furthermore, in the KTR cohort, multiple enzymes in this metabolic pathway were dysregulated during the recovery phase and showed differential expression between the two patient communities (**Supplementary Figure 1j-q**).

Following FA-AKI, there was a marked increase in metabolites related to arginine biosynthesis, with citrulline among the top metabolites significantly elevated (**Figure 4e, Supplementary Data File**). CB1R blockade effectively reduced the downstream metabolites, restoring aspartate, arginosuccinate, and arginine to physiologic levels (**Figure 4e**). This reduction resulted from decreased expression of arginosuccinate synthase (ASS1), which converts citrulline to arginosuccinate, and was mitigated by CB1R inhibition (**Figure 4f, g**). Post-AKI also increased arginase (ARG2) expression, leading to elevated levels of ornithine and proline (**Figure 4e**, **f, h**). JD5037 decreased ARG2 expression and the abundance of its downstream metabolites (**Figure 4e, f, h**). The L-arginine:glycine amidinotransferase (GATM) pathway was altered post-AKI, with reduced glycine and guanidinoacetate levels and decreased enzyme expression (**Figure 4e, f, i**), though not affected by JD5037, suggesting minimal involvement in this pathway. Notably, the metabolites mostly affected by JD5037 treatment were those whose abundance was increased at 14, and not 2 days, post-FA injection (**Supplementary Figure 7d, e**).

Furthermore, to specifically explore the effect of CB1R antagonism on arginine metabolism, healthy mice were treated with JD5037 for 1 week, revealing reductions in kidney arginine metabolites similar to those observed in vehicle-treated controls (**Supplementary Figure 8a**). Serum precursors for renal arginine synthesis, citrulline and arginosuccinate, were also decreased with JD5037 treatment, while systemic arginine levels remained unchanged (**Supplementary Figure 8b**). No changes in arginine-related metabolites were observed in the liver with JD5037 (**Supplementary Figure 8c**), underscoring kidney-specific effects. Altogether, peripheral CB1R blockade attenuated the profound alterations in renal arginine synthesis and metabolism during maladaptive repair, highlighting a novel therapeutic link between CB1R signaling and this metabolic pathway.

### Peripheral CB1R blockade suppresses the AKT-NF-κB inflammatory axis in maladaptive repair

Building on the observed normalization of metabolic and pro-fibrotic pathways by peripheral CB1R blockade, we next examined whether JD5037 modulates known CB1R-dependent intracellular signaling pathways that may contribute to the transition from injury to repair^32–34^. Post FA-AKI, molecular analysis revealed robust activation of several established cell-survival and inflammatory pathways, including mechanistic target of rapamycin complex 1 (mTORC1; as indicated by phosphorylated ribosomal protein S6 [p-S6]), extracellular signal-regulated kinase (ERK), and protein kinase B (AKT) (**Figure 5a-c, Supplementary Figure 4h-k**), reflecting broad pro-inflammatory and pro-remodeling signaling^35–38^. Notably, JD5037 treatment specifically reduced both phosphorylated and total AKT levels, without affecting the persistent activation of mTORC1 or ERK (**Figure 5a-c, Supplementary Figure 4h-k**), suggesting selectivity in CB1R-driven signaling.

**Figure 5.**
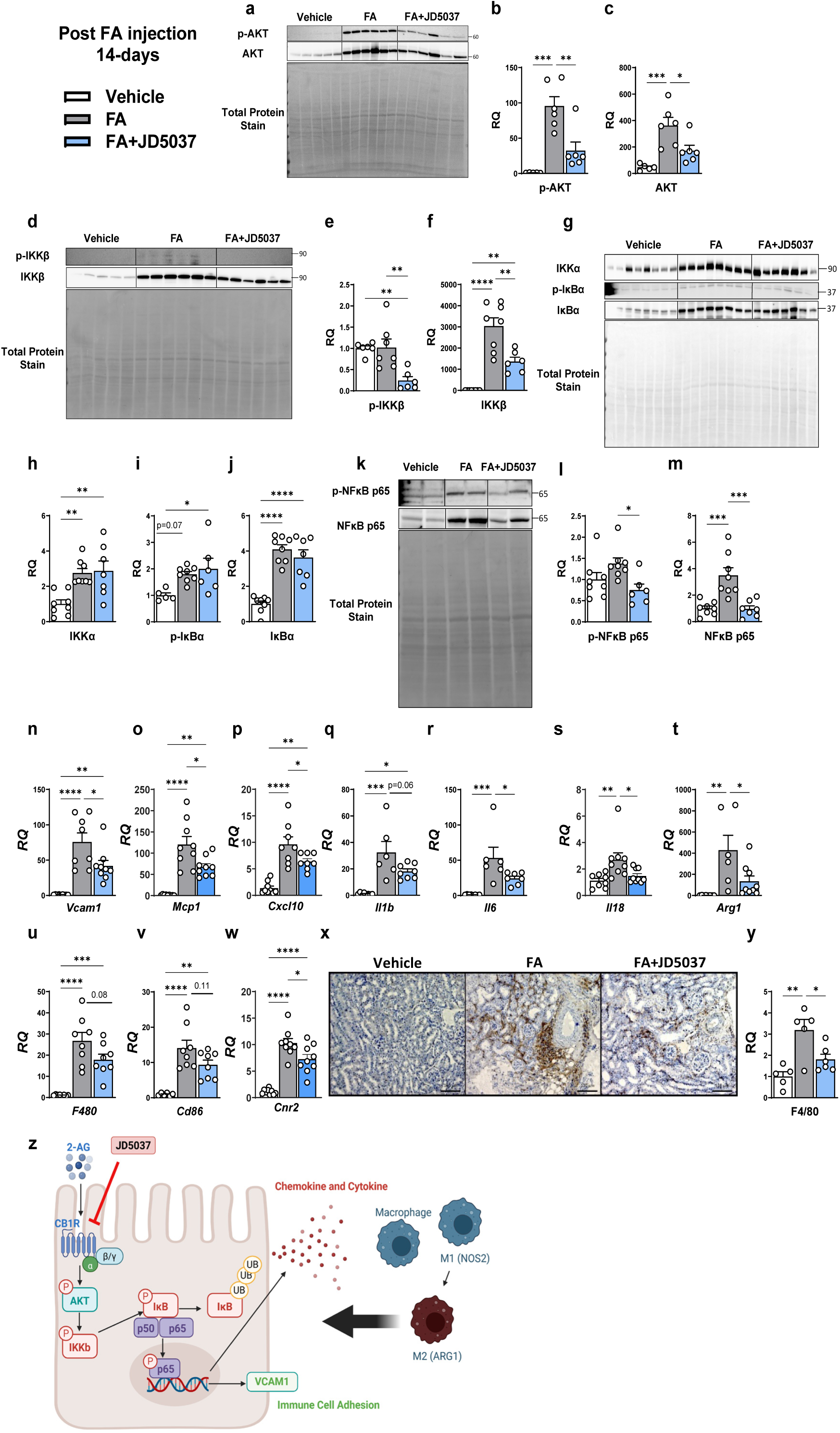
Peripheral CB1R blockade reduces NF-κB signaling and inflammatory response after AKI. FA-induced AKI (240 mg/kg) was induced in mice treated daily with the peripherally restricted CB1R inverse agonist, JD5037 (3 mg/kg) or vehicle. **(a)** Immunoblot analysis of phosphorylated and total AKT, and **(b-c)** their quantification (normalized to total protein). **(d)** Immunoblot for phosphorylated and total IKKβ, **(e-f)** with quantification (normalized to total protein). **(g)** Immunoblot for IKKα, phosphorylated and total IκBα, **(h-j)** with quantification (normalized to total protein). **(k)** Immunoblot for phosphorylated and total NF-κB p65 and **(l-m)** quantification (normalized to total protein). **(n-s)** Renal transcript levels for NF-κB-regulated inflammatory genes: *Vcam1*, *Mcp1*, *Cxcl10*, *Il1b*, *Il6*, *Il18.* **(t-w)** Transcript expression for immune and macrophage markers: *Arg1*, *F480 (Adgre1)*, *Cd86*, and *Cnr2*. **(x, y)** Representative kidney immunohistochemistry staining and quantification for F4/80 expression of infiltrating macrophages. Scale bar: 100 μm, magnification 20×. **(z)** Schematic illustration of the cellular signaling pathway modulated by peripheral CB1R blockade. Data represent the mean ± SEM (n = 5-9 per group). One-way ANOVA with Tukey’s post-hoc test was used for comparisons. *p < 0.05, **p < 0.01, ***p < 0.001, ****p < 0.0001. *Abbreviations:* CB1R, cannabinoid-1 receptor; FA, folic acid; AKI, acute kidney injury; NF-κB, nuclear factor-κB; AKT, protein kinase B; IKKα/β, inhibitor of κB kinase α/β; IκBα, inhibitor of κB alpha; *Vcam1*, vascular cell adhesion molecule 1; *Mcp1*, monocyte chemoattractant protein-1; *Cxcl10*, C-X-C motif chemokine ligand 10; *Il1b*, interleukin-1β; *Il6*, interleukin-6; *Il18*, interleukin-18; *Arg1*, arginase-1; *F4/80*, *Adgre1*-adhesion G protein-coupled receptor E1; *Cd86*, CD86 antigen; *Cnr2*, cannabinoid receptor-2

Because AKT can directly modulate nuclear factor-κB (NF-κB)^39^, a central transcriptional driver in post-AKI inflammation, immune recruitment, and failed renal repair^8,28,40^, we next assessed whether JD5037 alters this pathway. AKT regulates NF-κB activity in part through the inhibitor of κB kinase (IKK) complex. At 14-days following FA-AKI, expression of both IKKα and IKKβ increased substantially, and JD5037 reduced not only phosphorylated (active) IKKβ but also total IKKβ protein levels (**Figure 5d-h**). Consistent with enhanced NF-κB signaling, both phosphorylated and total inhibitor of κB alpha (IκBα), the canonical cytoplasmic NF-κB inhibitor, were markedly upregulated post-AKI; however, JD5037 did not affect IκBα expression or phosphorylation (**Figure 5g, i-j**). In contrast, total and phosphorylated levels of NF-κB p65, the transcriptionally active complex subunit, were strongly elevated post-AKI and were robustly suppressed by CB1R blockade (**Figure 5k-m**).

In line with reduced NF-κB activity, JD5037 treatment significantly lowered mRNA levels of *Vcam1* (a failed-repair PTC marker), chemokines (*Mcp1*, *Cxcl10*), and pro-inflammatory cytokines (*Il1b*, *Il6*, *Il18*), all of which are canonical NF-κB target genes (**Figure 5n-s**). Notably, most of these inflammatory mediators were predominantly elevated at later disease stages (**Supplemental Figure 2o-q**), consistent with the timing of CB1R upregulation and JD5037’s effects.

Given the observed reduction in inflammatory gene expression, we next evaluated whether CB1R blockade modulates immune cell recruitment, particularly macrophage infiltration and activation. This process is central to the maladaptive response^41–45^ and was also evident in response to injury in the various surgical AKI models. Specifically, expression of macrophage-associated markers, including F4/80 (*Adgre1*)*, Cd68, Cd86, Cd206* (*Mrc1*)*, Il1b, Arg1, Il18rap,* and *Il18r1*, was observed during the maladaptive phase following uIRI and bIRI (**Supplementary Figures 6 and 9**). In the FA-AKI model, JD5037 treatment significantly reduced renal transcriptional expression of arginase 1 (*Arg1*), a marker of alternatively activated (M2-type) macrophages, and showed a trend toward reduced expression of F4/80 (*Adgre1*) and CD86 (*Cd86*), markers of total and pro-inflammatory (M1-type) macrophages, respectively (**Figure 5t-v**). This was further supported by a reduction in *Cnr2* transcript levels (**Figure 5w**), enriched in inflammatory cells^46^, while JD5037 did not significantly alter *Cnr1* expression or eCB levels at this stage (**Supplementary Figure 4l-n**). Consistent with these findings, JD5037 treatment significantly decreased renal F4/80 expression (**Figure 5x, y**), indicating reduced macrophage infiltration. Collectively, these results demonstrate that peripheral CB1R blockade disrupts the AKT-NF-κB signaling axis and suppresses downstream inflammatory gene activation and immune cell recruitment (**Figure 5z**), thereby directly linking CB1R antagonism to mitigation of the maladaptive post-AKI inflammatory response and tubular injury.

### CB1R activation directly mediates the failed repair of proximal tubular cells

To validate the direct role of CB1R in mediating FR-PTCs, we next evaluated the effect of elevated 2-AG levels on *VCAM1* expression in primary human kidney proximal tubular cells (hKPTCs) by treating cells with the synthetic 2-AG analog, noladine ether (NE), or the MGLL inhibitor JZL184, both of which increased *VCAM1* expression levels (**Figure 6a**), directly connecting 2-AG-induced CB1R activation to FR-PTCs. Next, we treated hKPTCs with TNFα and NE to mimic the proinflammatory and elevated endocannabinoid ‘tone’ observed during the AKI-to-CKD transition in the presence/absence of JD5037. Indeed, the combined treatment significantly enhanced the transcriptional levels of failed-repair adhesion molecules and chemokine (**Figure 6b-d**), proinflammatory cytokines (**Figure 6e, f**), and arginine metabolism-related enzymes (**Figure 6g, h**). Crucially, CB1R antagonism significantly reduced these elevations, indicating a direct role of CB1R in mediating failed repair, pro-inflammatory responses, and metabolic dysregulation in the proximal tubules.

**Figure 6.**
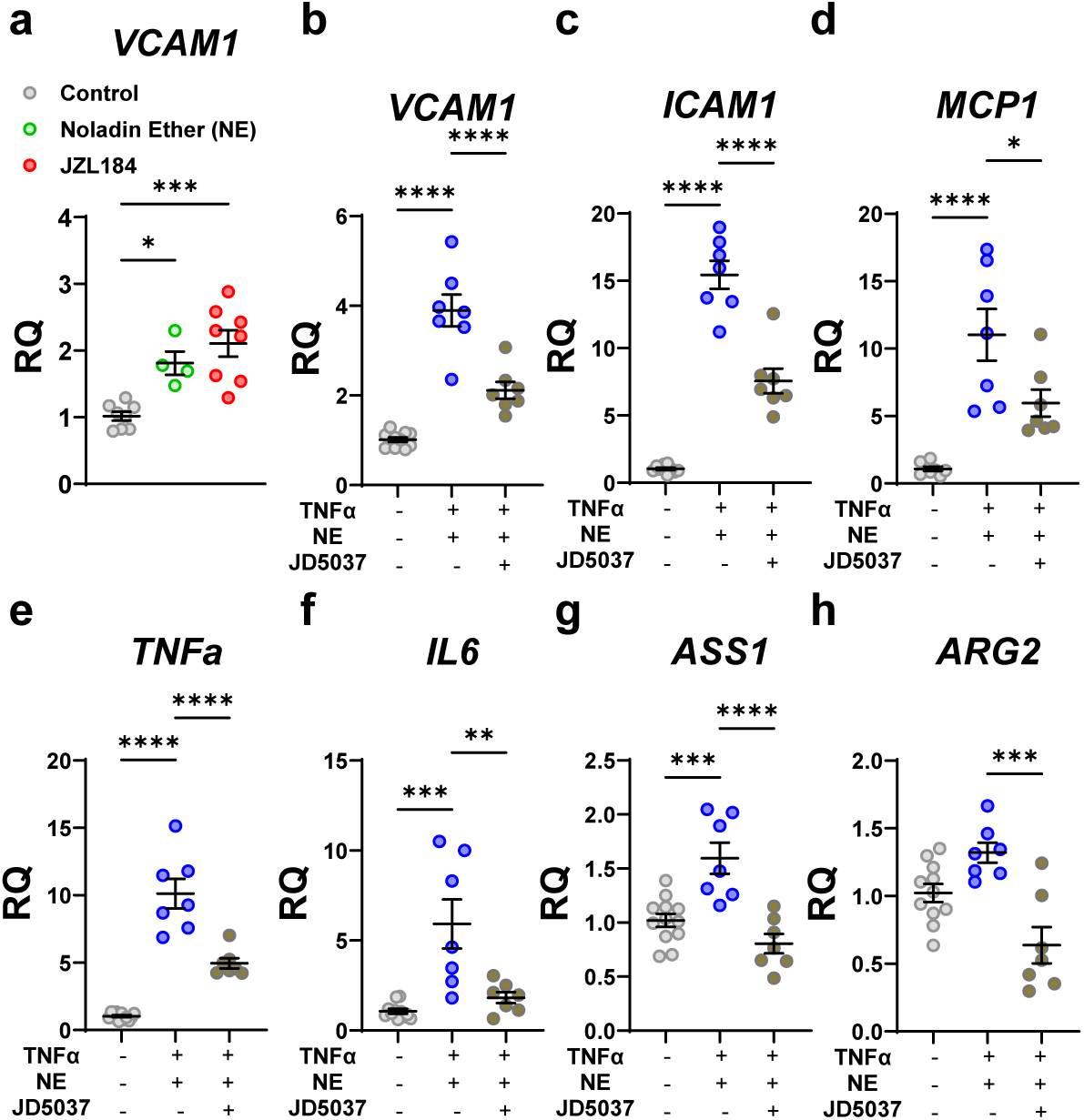
CB1R activation mediates failed repair in proximal tubular cells. Primary human kidney proximal tubular cells (hKPTCs) were plated in REGM. After 18 h of serum starvation, hKPTCs were treated with vehicle (DMSO), noladin ether (NE; 1 µM), or JZL184 (1 µM) for 6 h or pre-incubated for 1 h with JD5037 (100 nM) and then treated with TNFα (20 ng/mL) and NE (1 µM) combination. After 6 h, cells were collected for mRNA extraction and qPCR. The transcriptional levels of (**a**) *VCAM1* after CB1R activation with NE or JZL184. The transcriptional levels of (**b**) *VCAM1*, (**c**) *ICAM1*, (**d**) *MCP1*, (**e**) *TNFa*, (f) *IL6*, (**g**) *ASS1*, and (**h**) *ARG2* after the combined treatment with TNFα and NE, in the absence or presence of JD5037. Data represent the mean ± SEM (n = 4-12 per group). One-way ANOVA with Tukey’s post-hoc test was used for comparisons. *p < 0.05, **p < 0.01, ***p < 0.001, ****p < 0.0001. *Abbreviations: VCAM1*, vascular cell adhesion molecule 1; *ICAM1*, Intercellular Adhesion Molecule 1; *MCP1*, Monocyte Chemoattractant Protein-1; *TNFa*, Tumor Necrosis Factor alpha; *IL6*, interleukin-6; *ASS1*, argininosuccinate synthase 1; *ARG2*, arginase-2.

## Discussion

Effective management strategies for AKI and its progression to CKD are lacking, underscoring the imperative to define the molecular determinants of maladaptive versus successful repair. Across both human and murine models, our study establishes that CB1R expression and signaling are selectively increased during late-stage kidney injury, specifically in PTCs that fail to repair. This temporal and cellular specificity links CB1R activity to maladaptive kidney remodeling, suggesting a critical role for phase-specific intervention during the AKI-to-CKD transition.

Our data demonstrate that the maladaptive phase after AKI is characterized by upregulation of key ECS components, notably CB1R, its ligand 2-AG, and the biosynthetic enzyme DAGL. While injury etiology may influence initial molecular events^47^, the convergence on CB1R upregulation and elevation of its endogenous ligand is a hallmark of failed repair, which supports clinical reports showing that CB1R overactivation correlates with worse kidney function^12,48^, and is upregulated in patients with chronic allograft dysfunction^26^ and non-metabolic CKD^14^. Single-nucleus RNA sequencing confirms that CB1R elevation occurs specifically in the subset of PTCs, termed FR-PTCs, whose proinflammatory, profibrotic gene signature likely drives CKD progression^7,25,28,40^, and this study further demonstrates that 2-AG-induced CB1R activation promotes a failed-repair phenotype in KPTCs. Furthermore, peripheral CB1R blockade during this maladaptive phase ameliorates tubular injury, inflammation, and metabolic perturbations. The less pronounced effect of CB1R antagonism on fibrosis in our FA-AKI model may reflect a limited treatment window, with fibrosis peaking later than the intervention period; this contrasts with models such as UUO, where CB1R inhibition robustly reduces fibrosis^14^.

Consistent with recent insights into metabolic rewiring during kidney injury and repair^49^, our study demonstrates that the ECS undergoes AKI stage-specific alterations. Importantly, although previous work suggested potential heterogeneity of renal eCBs^13^, their precise intrarenal distribution had not been confirmed. Using high-resolution spatial molecular imaging, we now provide direct evidence of the compartment-specific localization of eCBs in both healthy and injured kidneys. This approach further reveals a dynamic temporal shift in the distribution of 2-AG and AEA across disease stages, underscoring the spatial-metabolic complexity of ECS remodeling during AKI.

Although the pathogenic involvement of the ECS, and particularly CB1R, is well-demonstrated in diabetic and other chronic kidney conditions^15–19^, its acute-phase role in AKI has been controversial. Our findings indicate that CB1R expression remains low during acute insult and that CB1R blockade does not mitigate early damage, contradicting findings in cisplatin^20^ and gentamicin^50^-induced nephropathy. These differences likely reflect the distinct mechanisms underlying each type of renal pathology, indicating varying ECS effects depending on AKI etiology. Further investigation of the effect of direct CB1R activation during the acute stage is needed, as CB1R agonists or eCB elevation have been shown to protect against AKI^23,51–53^. Importantly, CB1R-independent pathways may also be involved in this stage, as the protective effect of FAAH inactivation after IRI was due to specific AEA metabolites rather than direct CB1R activation^54^. In line with this, our group found elevated systemic AEA levels in non-obese patients after partial nephrectomy, suggesting eCBs respond to acute renal ischemia^48^. Additionally, many studies show CB2R regulation in AKI and fibrogenesis, though its distribution in the kidney^55^ and protective^14,51,56,57^ or deleterious^58–60^ role is highly debated. Nevertheless, our data across human and animal systems reinforce the mechanistic primacy of CB1R in the chronic, not acute, phase of kidney injury.

Arginine metabolism plays a central role in normal renal physiology, contributing to processes such as nitric oxide production, ammonia handling, and endogenous arginine and creatine synthesis^30,61,62^. Studies focusing on acute renal insult have shown that this metabolic network becomes markedly dysregulated, specifically with ARG2 upregulation, aggravating endothelial dysfunction, oxidative stress, and tubular injury^63–66^. Our untargeted metabolomics showed excessive arginine turnover during the maladaptive repair phase, corroborating its known role in cell division, healing, and inflammation^67^. Consistent with this, arginine-pathway metabolites accumulated at 14 days but not 2 days post-FA-AKI, and KTRs with maladaptive repair showed altered expression of enzymes governing arginine metabolism. CB1R antagonism normalized the pathological accumulation of arginine and its metabolites and mitigated disruption of key enzymes involved in its turnover (ASS1, ARG2), both *in-vivo* and hKPTCs. Indeed, citrulline accumulation and disruption of arginine metabolic enzymes have been identified as a key feature of CKD in other renal injury models^63,68–75^. The interplay between CB1R signaling and NO synthesis^76,77^ further supports a novel mechanistic link to arginine metabolism. Future research is needed to define cell- and context-specific regulatory mechanisms as both the receptor and metabolic enzyme effects vary by cell type and tissue^14,78–80^.

Mechanistically, the benefit of peripheral CB1R blockade extends to suppression of the AKT-NF-κB signaling axis. While several reports have shown that transient AKT activation may be protective during acute injury^81,82^, sustained activation promotes maladaptive repair and fibrosis^83,84^. By disrupting AKT activation, CB1R antagonism reduces downstream IKKβ phosphorylation, blunts NF-κB transcriptional activity, and downregulates proinflammatory and profibrotic gene programs. Indeed, hyperactive NF-κB signaling has long been implicated in renal inflammation and fibrosis^8^, and its contribution to failed-repair phenotype is supported by increased chromatin accessibility for NF-κB binding motifs^7^ as well as activation of NF-κB target genes in these FR-PTC^28^. In KTRs, NF-κB complex proteins and target genes were significantly elevated in chronically injured cells, further supporting their clinical relevance in AKI-to-CKD progression^28^. It is increasingly clear that CB1R-NF-κB interactions are cell- and context-dependent^86–88^; further studies are needed to map these in the post-AKI setting and to identify optimal timing and targets for intervention.

In summary, this study highlights the distinct phenotypic, metabolic, and ECS changes during kidney injury and repair, with CB1R overexpression and overactivation emerging in the later stage. We show that peripheral CB1R blockade does not affect AKI development but reduces renal damage, inflammation, and metabolic disturbances as the disease progresses. These findings underscore the importance of time-dependent changes in ECS-targeting for AKI treatment and the therapeutic potential of peripheral CB1R blockade in mitigating maladaptive kidney repair.

## Data Sharing Statement

The authors confirm that the data supporting the findings of this study are available in the article and its Supplementary Material at www.kidney-international.org.

## Acknowledgments

We would like to acknowledge Yael Calles for her technical assistance. We would also like to acknowledge Prof. Dr. Pietro Cippà of the University of Zurich for providing clinical data for the KTR study, as well as Dr. Yuval Nevo and Dr. Sharona Elgavish from the Azrieli Omics Center for their assistance with the bioinformatics analysis.

## Author Contributions

***Conceptualization:*** Ariel Rothner, Liad Hinden, Joseph Tam.

***Funding acquisition:*** Joseph Tam.

***Investigation:*** Ariel Rothner, Liad Hinden, Aviram Kogot-Levin, Shridhar Betkar, Elisheva Benkovitz, Amani Zoabi, Anna Permyakova, Asaf Kleiner, Vladislav Nesterenko, Alina Nemirovski, Ifat Abramovich, Bella Agranovich, Inbar Plaschkes.

***Resources:*** Joseph Tam, Eyal Gottlieb, Gil Leibowitz.

***Supervision:*** Joseph Tam.

***Validation:*** Joseph Tam.

***Writing – original draft:*** Ariel Rothner.

***Writing – review & editing:*** Liad Hinden, Joseph Tam.

