## Supplemental Information, Figures, and Tables for "Peripheral CB1R Blockade Suppresses AKI-to-CKD Maladaptive Repair"

### Inventory of Supporting Information

- **Supplemental Methods and References**
- **Supplementary Figure 1.** Gene expression disruptions in the kidneys of KTRs at acute and chronic disease stages.
- **Supplementary Figure 2.** FA-induced AKI and maladaptive repair stages display distinct phenotypic signatures
- **Supplementary Figure 3.** Peripheral CB1R blockade does not alter early FA-induced AKI severity
- **Supplementary Figure 4.** Peripheral CB1R blockade post FA-AKI does not affect renal glucose handling and signaling pathways
- **Supplementary Figure 5.** ECS changes in rodent models of AKI and maladaptive repair.
- **Supplementary Figure 6.** Enhanced CB1R expression and activation in murine IRI models.

- **Supplementary Figure 7.** Differential changes in arginine-related metabolites in FA- AKI and maladaptive repair.
- **Supplementary Figure 8.** Peripheral CB1R blockade in healthy mice modulates arginine-related metabolites in serum and kidney, but not liver.
- **Supplementary Figure 9.** Enhanced macrophage expression and activation in post-AKI maladaptive repair.
- **Supplementary Table 1.** Endocannabinoid analysis following FA-induced AKI
- **Supplementary Table 2.** LC-MS/MS instrument settings for endocannabinoid analysis.
- **Supplemental Table 3.** Mouse primer sequences used for qPCR.
- **Supplemental Table 4.** Human primer sequences used for qPCR.
- **Supplementary Table 5.** List of antibodies used in the study.
- **Supplemental Data 1.** Metabolomics analysis at 2 and 14-days following FA-induced AKI compared to vehicle control.
- **Supplemental Data 2.** Kidney metabolomics results for vehicle vs. 2-days post FA-injection
- **Supplemental Data 3.** Kidney metabolomics results for 14-days post FA-injection: Vehicle vs FA vs FA+JD5037

**Supplementary Methods**

***Transcriptomics Analysis***

*Clinical Dataset:* The clinical dataset of kidney transplant recipients (KTRs) bulk RNA-seq (GSE162805)[1] was also obtained from the GEO database. This dataset comprises protocol biopsies from 42 KTRs at four time points related to kidney transplantation: before implantation (PRE, kidney flushed and stored on ice), after reperfusion (POST, at the end of the surgical procedure), and at 3- and 12-month post-implantation (follow-up). Genome-wide gene expression profiling using RNA-Seq was conducted as previously described[1].

Briefly, samples from the 3- and 12-month follow-ups were clustered into densely connected networks (communities) using the Walktrap community detection algorithm. Community A displayed a transcriptional signature associated with kidney injury, fibrosis, and chronic inflammation, while Community B represented healthier counterparts. These communities were clinically relevant, as Community A exhibited worse kidney function [measured by estimated glomerular filtration (eGFR)] and a higher fibrosis score[1].

The raw counts matrix was downloaded from GEO. Genes with a sum of counts lower than 20 across all samples were filtered out. Normalization and differential expression analysis were conducted using the DESeq2 package with default parameters, except not using the independent filtering algorithm. Differentially expressed genes (DEGs) were detected between the following sample categories: before and immediately after reperfusion samples (Pre vs. Post), between follow-up time to immediately after reperfusion for each community (follow-ups vs. Post), and between communities (A vs. B) at follow-up time points. A significant DEG threshold was set at a false discovery rate (FDR)-adjusted p-value $< 0.1$.

*Mouse Datasets:* Mouse bulk RNA-seq (GSE98622)[2] was retrieved from the GEO database, covering multiple time points following bilateral renal IRI, including 2, 4, 24, 48, and 72 hours, as well as 7, 14, and 28 days, 6 months, and 12 months post-IRI, along with sham controls (4 hours, 24 hours, and 12 months). As reported, PCA analysis revealed similar expression patterns across sham surgery samples; all sham time points were combined to serve as a control for comparison[2]. A time-course analysis of key gene expression following IRI was performed.

Datasets of mouse kidneys subjected to bilateral ischemia-reperfusion injury (IRI) were accessed through the Kidney Interactive Transcriptomics (KIT) database (https://humphreyslab.com/SingleCell/). We accessed single-nucleus ATAC-seq (snATAC-seq) data from Muto *et al*[3] and single-nucleus RNA-seq (snRNA-seq) data from Kirita *et al.*[4]. As described, both studies profiled whole-kidney nuclei across multiple time points following bilateral IRI, enabling characterization of chromatin accessibility and transcriptional trajectories during injury, repair, and maladaptive repair. We directly queried gene expression and chromatin accessibility of Cnr1 across defined cell clusters and time points within KIT.

#### *Animals*

All experimental protocols were approved by the Institutional Animal Care and Use Committee of the Hebrew University of Jerusalem (AAALAC accreditation #1285; Ethic approval numbers MD-19-15935 and MD-22-17030) and comply with ARRIVE guidelines. Male C57BL/6J mice, aged 10-12 weeks, were obtained from Harlan (Israel). The mice were housed in a conventional animal facility under specific pathogen-free (SPF) conditions, with controlled temperatures of 22-24°C, 55 ± 5% humidity, and a 12-hour light/dark cycle. Food and water were provided *ad libitum*.

***Folic acid-induced acute kidney injury (FA-AKI)***

Mice were induced with acute kidney injury (AKI) through a single intraperitoneal (i.p.) injection of 240 mg/kg folic acid (FA, Cayman Chemicals) dissolved in a 0.3 M sodium bicarbonate solution or received a vehicle injection (injection volume of 10 µL/g, an average of 270 µL per mouse). Mice were monitored for body weight changes, morbidity, and mortality, and received daily i.p. injections of sterile 0.9% saline starting 2 days after the FA injection to prevent dehydration. Twenty-four-hour urine output was measured and collected at 2-, 14-, and 28-day post-FA injection using the CCS2000 Chiller System (Hatteras Instruments). Mice were then euthanized by cervical dislocation under anesthesia; trunk blood, kidneys, and liver were collected, with samples either snap-frozen or fixed in buffered 4% formalin for further analysis.

To assess the effect of peripherally restricted CB1R blockade on the kidney, a separate cohort of male C57Bl/6J mice was induced with FA-AKI or sodium bicarbonate vehicle. These mice were treated daily with JD5037 (Haoyuan Chemexpress Co., Ltd, 3 mg/kg, p.o.) or placebo (1% Tween80, 4% DMSO, 95% Saline) for 2- or 14-day following FA-injection. The effect of JD5037 treatment was also assessed in a healthy cohort, where mice were treated for 7 days with JD5037 (3 mg/kg, i.p.) or vehicle (1% Tween80, 4% DMSO, 95% Saline, i.p.). The animals were euthanized, and organs were collected as described above.

***Unilateral ischemia-reperfusion injury with delayed nephrectomy (uIRI)***

C57Bl/6J mice, 12-13 weeks old, male mice underwent renal pedicle clamping for 21 min to induce IRI to the left kidney on day 0, followed by a right kidney nephrectomy on day 14. Sham control mice underwent an identical surgical procedure, with the exception of ischemic clamping. Mice were monitored daily for body weight reduction, morbidity, and mortality. Long-lasting buprenorphine (3.5 mg/kg) was administered by SC injection for analgesia on days 0 and 14 before the surgeries. On days 3 and 15, mice were euthanized by cervical dislocation under anesthesia; trunk blood and kidneys were collected, and samples were snap-frozen for further analysis.

***Bilateral ischemia-reperfusion injury (bIRI)***

C57Bl/6J male mice, 12-13 weeks old, underwent renal pedicle clamping for 28 min on day 0 to induce IRI in both kidneys. Sham control mice underwent identical surgical procedures, with the exception of ischemic clamping. Mice were monitored daily for body weight reduction, morbidity, and mortality. Long-lasting buprenorphine (3.5 mg/kg) was administered by SC injection for analgesia on day 0 of surgery. On days 2 and 14, mice were euthanized by cervical dislocation under anesthesia; trunk blood and kidneys were collected, and samples were snap-frozen for further analysis.

The experimental unit in all *in vivo* studies was a single mouse. Animals were allocated to the relevant treatment and time-point groups as described above, and the exact number of mice per group is reported in each figure legend. Across all AKI mouse models as well as healthy control studies, the total number of animals used is fully specified in the figure legends and supplementary figure legends.

***Blood and urine biochemistry***

Serum levels of blood urea nitrogen (BUN), creatinine, and glucose, as well as urine levels of creatinine and glucose, were determined using the Cobas C-111 biochemistry analyzer (Roche, Switzerland). Serum levels of Lipocalin-2 (NGAL) were analyzed by ELISA assay (Abcam; Cat# ab199083) according to the manufacturer's instructions. Creatinine clearance rate (CrCl) was calculated using urine and serum creatinine levels *(CrCl [mL/h] = Urine creatinine [mg/dL] × Urine volume/Serum creatinine [mg/dL] × 24 h)*.

***LC-MS-based metabolomics***

Comparative kidney metabolomic analysis was performed at different points following FA-AKI (2 and 14 days) and for samples treated with a peripheral CB1R inverse agonist. Additional metabolite analysis was performed on kidney, liver, and serum samples in healthy mice treated with a peripheral CB1R inverse agonist. Each comparison was made against the appropriate vehicle-treated controls. Serum samples were diluted 1:10 with an extraction solution (75:25 methanol: acetonitrile). Frozen tissue samples were transferred into soft tissue homogenizing tubes containing 1.4 mm ceramic beads (CK14, Bertin Corp., Rockville, MD, USA), prefilled with 400 µL of cold (−20 °C) metabolite extraction solvent (methanol:acetonitrile:water, 5:3:2), and kept on ice. Samples were homogenized using a Precellys 24 tissue homogenizer cooled to 4 °C (3 × 20 s at 6,000 rpm, with a 30 s gap between cycles; Bertin Technologies, Montigny-le-Bretonneux, France). Homogenized extracts were centrifuged at 18,000 × g for 15 min at 4 °C; the supernatants were collected in microcentrifuge tubes and centrifuged again at 18,000 × g for 10 min at 4 °C. The supernatants were transferred to glass HPLC vials and kept at −75 °C prior to LC-MS analysis.

LC-MS metabolomic analysis was performed as described previously[5]. Briefly, a Dionex Ultimate 3000 high-performance liquid chromatography (UPLC) system coupled to an Orbitrap Q-Exactive Mass Spectrometer (Thermo Fisher Scientific) was used with a resolution of 35,000 at 200 mass/charge ratio (m/z), electrospray ionization, and polarity switching mode to enable both positive and negative ions across a mass range of 67 to 1000 m/z. The UPLC setup consisted of a ZIC-pHILIC column (SeQuant; 150 mm × 2.1 mm, 5 µm; Merck, Darmstadt, Germany) with a ZIC-pHILIC guard column (SeQuant; 20 mm × 2.1 mm). A total of 5 µL of the extracts was injected, and the compounds were separated with a mobile phase gradient of 15 min, starting at 20% aqueous (20 mM ammonium carbonate adjusted to pH 9.2 with 0.1% of 25% ammonium hydroxide) and 80% organic (acetonitrile) and terminated with 20% acetonitrile. The flow rate and column temperature were maintained at 0.2 mL/min and 45 °C, respectively, for a total run time of 27 min. All metabolites were detected using mass accuracy below 5 ppm. Thermo Xcalibur was used for the data acquisition.

The analyses were performed with TraceFinder 5.0 (Thermo Fisher Scientific, Waltham, MA, USA), identifying the exact mass of the singly charged ion and known retention time, using an in-house MS library built by running commercial standards for all metabolites (550 metabolites). Data from each sample were normalized to its corresponding total ionization count (serum), tissue weight, or tissue weight × total ionization count. Metabolite-Auto Plotter 2 was used for data normalization and visualization during data processing[6]. Untargeted analysis was performed using the Compound Discoverer™ Software (ThermoFisher Scientific).

Statistical analysis was performed by MetaboAnalyst 5.0. Differentially abundant metabolites at each time point compared to their appropriate vehicle control were quantified ($p < 0.05$). Selecting only the metabolites detected in all groups (>250), metabolic enrichment analysis using log-transformed data, comparing each disease state (2 days or 14 days) to their respective pre-treatment group (0 days), was performed using MetaboAnalyst 5.0, utilizing the small molecule pathway database (SMDPBP) metabolic pathway library, including only pathways identifying >2 metabolites.

***Histopathology***

Periodic acid-Schiff (PAS) staining (Abcam; Cat# ab150680) was performed on paraffin-embedded kidney sections (4 µm) to assess tubular injury in a blinded manner. Twelve random images from each kidney were captured using an AxioCam ICc5 color camera mounted on an Axio Scope.A1 light microscope (Zeiss, Germany). Tubular injury was analyzed using Image Pro Plus 6.0 software, with injury characterized by tubular necrosis, cast formation, tubular dilation, and brush border loss. The area of injury was quantified and calculated as a percentage of the total tissue area.

Trichrome staining (Abcam; Cat# ab150686) was performed on paraffin-embedded kidney sections. Images were captured as described above, and quantification was performed in Photoshop, with a minimum of 12 random kidney sections per mouse. The relative fold-change in collagen staining compared to the control was calculated. The figures show the animal with the median value for each group.

#### *Immunohistochemistry*

Kidney tissues were fixed, paraffin-embedded, and sectioned at 4 µm thickness. Sections were deparaffinized, rehydrated through graded alcohols, and subjected to heat-induced epitope retrieval in 10 mM citrate buffer (pH 6.0; Thermo Scientific, IL, USA). To quench endogenous peroxide activity, sections were incubated in 3 % hydrogen peroxide ($H_2O_2$) in methanol for 10 minutes at room temperature. Non-specific binding was blocked by incubating the sections in 2.5 % horse serum (Cat# VE-S-2000, Vector Laboratories) for 1 hour at room temperature. Slides were then incubated overnight at 4 °C with a F4/80 (D2S9R) Rabbit monoclonal antibody (Cat# 70076, Cell Signaling; 1:300 dilution). After rinsing, sections were incubated with ImmPRESS-HRP horse anti-rabbit secondary antibody (Cat# VE-MP-7401, Vector Laboratories) for 1 hour at room temperature. An immunoreactive signal was developed using 3,3'-diaminobenzidine (DAB) substrate (Cat# SK-4105, Vector Laboratories, USA) until the desired staining intensity was achieved. Slides were counterstained with hematoxylin, dehydrated, cleared, and mounted using Vecmount mounting medium (Cat# H-5000, Vector Laboratories, USA). Stained sections were visualized by light microscopy, and F4/80 expression was quantified using ImageJ software. A consistent color threshold was applied to define DAB-positive areas. For each sample, at least 5 random, non-overlapping fields were analyzed to ensure robust, reproducible quantification across biological replicates.

#### *Endocannabinoid analysis*

Extraction, purification, and quantification of kidney and urine endocannabinoids (eCBs) were performed using stable isotope dilution liquid chromatography/tandem mass spectrometry (LC-MS/MS) as previously described[7,8]. In brief, kidney samples were homogenized in ice-cold Tris Buffer, and protein concentration was determined by the BCA assay. For urine samples, proteins were first precipitated with ice-cold acetone and Tris buffer (50 mM, pH 8.0). For both mediums, an ice-cold extraction buffer (1:1 MeOH/Tris Buffer + internal standard [$d_4$-AEA and $d_5$-AG]) was added. Homogenates were then extracted using a mixture of ice-cold $CHCl_3$:MeOH (2:1, vol/vol), followed by three washes with ice-cold chloroform. Samples were dried under a nitrogen stream and reconstituted in MeOH.

Analysis was performed using an AB Sciex (Framingham, MA, USA) QTRAP® 6500+ mass spectrometer coupled with a Shimadzu (Kyoto, Japan) UHPLC System. Chromatographic separation was achieved using a Kinetex 2.6 µm C18 (100 × 2.1 mm) column from Phenomenex (Torrance, CA, USA). The autosampler temperature was set at 4 °C, and the column was maintained at 40 °C. Gradient elution mobile phases consisted of 0.1% formic acid in water (phase A) and 0.1% formic acid in acetonitrile (phase B).

eCBs were detected in positive-ion mode by electron spray ionization (ESI) and multiple reaction monitoring (MRM). The collision energy (CE), declustering potential (DP), and collision cell exit potential (CXP) for the monitored transitions are provided in **Supplementary Table 2**. Levels of anandamide (AEA) and 2-arachidonoylglycerol (2-AG) in samples were measured against standard curves and calculated in pmol/mg kidney weight, pmol/mg protein, or pmol/hour as indicated. For time-course analysis, the fold change of each eCB relative to pre-FA injection levels was calculated across three separate studies.

***Desorption electrospray ionization spectrometry (DESI-MSI)***

Frozen kidney tissues were sectioned at 20 μm thickness using a cryostat microtome CM1950 (Leica, Germany). Inert embedding media and optimal cutting temperature (OCT) were used to adhere the sample to the specimen mount, ensuring proper orientation for sectioning. Sections of kidneys from different time-points after FA-AKI were mounted on the same glass slide and scanned simultaneously by DESI-MSI. Prior to scanning, tissues were thawed for 10 minutes in a vacuum desiccator.

Kidney tissue sections were analyzed using Xevo G2-XS Q-TOF mass spectrometer (Waters Corp., USA) equipped with a 2D moving stage, achieving a spatial resolution of 50 μm. The mass spectrometer's heated ion transfer line was set to 30 °C. Mass spectra were acquired in positive-ion mode over the mass range 50-1200 m/z, with a resolving power of 30,000 FWHM and a spray voltage of 0.7 kV. The spray solvent consisted of methanol:water:formic acid (98:2:0.01, v/v/v), with a spray flow rate maintained at 2 μL/min. 2-AG and AEA were detected as $M^{+}Na^{+}$ at *m/z* 401.27 and 370.27, respectively

Data were processed using HDI V1.6 software (Waters, USA), which includes tools for unique ion imaging comparison and region-of-interest (ROI) analysis. Peak intensities in the analyte table and the MS spectrum were updated to represent data specific to selected *m/z* values within a given ROI. The cortex and medulla regions were selected as separate ROIs, and the average intensity for 2-AG or AEA ions was calculated for each diseased stage. Data were normalized by the average of each ion for each DESI-MSI scan.

***Fatty acid amide hydrolase activity***

Fatty acid amide hydrolase (FAAH) activity in kidney lysates was assessed using the FAAH Activity Assay Kit (Abcam; Cat# ab252895) following the manufacturer's instructions.

***Cell culture***

Primary human kidney proximal tubular cells (hKPTCs, Lonza; Cat #CC-2253) were cultured in REGM BulletKit medium (Lonza; Cat #CC-3191 & CC-4127). To test the effect of CB1R activation on failed repair, hKPTCs (passage 9, 0.8-1×$10^6$ cells per plate) were seeded in 10-cm culture dishes and allowed to adhere for 24 hours prior to experimentation. Following attachment, cells were serum-starved in serum-free medium (SFM) supplemented with 0.1% BSA for approximately 18 hours before treatment initiation. For CB1R activation, vehicle (DMSO), noladine ether (NE, 1 µM), or JZL184 (1 µM) was supplemented to the SFM for 6 hours. For CB1R blockade, cells were pretreated with JD5037 (100 nM) for 1 hour before exposure to TNFα (20 ng/mL), NE (1 µM), or their combination. After 6 hours of treatment, cells were collected for mRNA extraction and qPCR.

***Real-time PCR***

mRNA from the mouse kidney cortex or hKPTCs was extracted using Bio-Tri RNA lysis buffer (Bio-Lab, Israel) and treated with DNase I (Thermo Scientific, IL, USA). The RNA was reverse transcribed into cDNA using the qScript cDNA Synthesis kit (Quantabio). Real-time PCR was performed with iTaq Universal SYBR Green Supermix (Bio-Rad, CA) on a CFX Connect ST system (Bio-Rad, CA). The primers used for detecting mouse and human genes are listed in **Supplementary Tables 3 and 4**. Gene expression levels were normalized to *ribosomal protein S16*, Gapdh, or *RPLP* for mouse and human genes, respectively.

#### *Western Blotting*

Kidney homogenates were prepared in RIPA buffer (25 mM Tris-HCl, pH 7.6, 150 mM NaCl, 1% NP-40, 1% sodium deoxycholate, 0.1% SDS) using the BulletBlender® with zirconium oxide beads (Next Advanced, Inc., NY, USA). Protein concentrations were measured with the Pierce™ BCA Protein Assay Kit (Thermo Scientific, IL, USA). Samples were resolved by SDS-PAGE (4–15% acrylamide, 150 V) and transferred to PVDF or nitrocellulose membranes using the Trans-Blot® Turbo™ Transfer System (Bio-Rad, CA).

Membranes were blocked for 1 hour in 5% milk (in 1× TBS-T) to prevent nonspecific binding, then incubated overnight at 4 °C with primary antibodies (**Supplementary Table 5**). Horseradish peroxidase (HRP)-conjugated secondary antibodies were applied for 1 hour at room temperature. Chemiluminescence detection was performed using Clarity™ Western ECL Blotting Substrate (Bio-Rad, CA) and imaged with ChemiDoc™ Touch Imaging System (Bio-Rad, CA). Densitometry analysis was carried out using Bio-Rad CFX Manager software. Quantification was normalized to Rabbit anti-VCP (Abcam; Cat# ab204290, 1:1000) or total protein by ponceau staining.

#### *Quantification of folic acid*

To validate FA-AKI induction, FA concentrations in kidney samples were quantified by LC-MS/MS. Kidneys were dissolved in a methanol:acetonitrile:water (5:3:2) solution, with Methotrexate Hydrate (Sigma; Cat# A6770) included as an internal standard. The samples were then centrifuged, and the supernatant was analyzed by liquid chromatography-electrospray ionization-tandem mass spectrometry (LC-ESI-MS/MS). LC-ESI-MS/MS measurements were carried out on Shimadzu (Kyoto, Japan) UHPLC System coupled with a Sciex (Framingham, MA, USA) QTRAP® 6500$^+$ mass spectrometer. Liquid chromatographic separation was obtained using 5 µL injections of all standard and sample solutions onto a Cortecs C18 2.7 µm (100×2.1 mm) column from Waters (Ireland). The mobile phase consisted of solvent A (water containing 0.1% formic acid) and solvent B (acetonitrile containing 0.1% formic acid). The autosampler was set to 4 °C, and the column was maintained at 40 °C during the entire analysis.

The mobile phase gradient (flow rate of 0.3 mL/min) was programmed as follows: 95% A (v/v) from 0 to 2.0 min, 95-80%A (v/v) from 2.0-2.5 min, 80% A (v/v) from 2.5-3.5 min, 80%-5% A (v/v) from 3.5-6.0 min, 5% A (v/v) from 6.0-9.5 min, 5-95% A (v/v) from 9.5-10.0 min. The total run time was 15 min.

For the mass spectrometric analysis, the IonDrive™ Turbo V source operating in negative mode, the following parameters were set: source temperature: 600 °C; Ion Spray voltage: -4500V; curtain gas: 30 psi; nebulizer gas (Gas 1): 45 psi; turbo heater gas (Gas 2):55 psi. Nitrogen was used as a nebulizer and collision gas. Instrument settings optimized for folic acid were: Molecular ion $[M-H]^-$ (*m/z*)- 440.1, Fragment (*m/z*)- 310.9, DP (volts)- -40, CE (volts)- -30, CXP (volts)- -12; and methotrexate were: Molecular ion $[M-H]^-$ (*m/z*)- 453.0, Fragment (*m/z*)- 324.0, DP (volts)- -55, CE (volts)- -28, CXP (volts)- -17. Acquisition was performed in multiple reaction monitoring (MRM) mode, and the Analyst 1.7.3 software (Sciex, USA) was used for data acquisition and processing.

***Statistical analysis***

Statistical analyses were conducted using GraphPad Prism version 10. Data are presented as mean ± standard error. Outliers were identified and removed using the robust ROUT (Robust regression and Outlier removal) method, with a false discovery rate (Q) set to 1%. Differences between two groups were assessed using a two-tailed Student's t-test for pre-clinical samples (unpaired or paired, as appropriate for comparison within the same sample). For clinical samples, the Mann-Whitney test was used. Comparisons across multiple groups were performed using one-way ANOVA, followed by Tukey's multiple comparisons test for three-group comparisons, and Dunnett's test for comparison to the pre-injection state (for groups >3). Pearson's correlation coefficient and univariate linear regression analysis were applied for correlation analyses. To predict binary outcomes of interstitial fibrosis (ci) at 12 months post-transplantation, a logistic regression model was trained, and a receiver operating characteristic (ROC) curve was generated, with the area under the ROC curve (AUROC) calculated. A p-value $< 0.05$ was considered statistically significant.

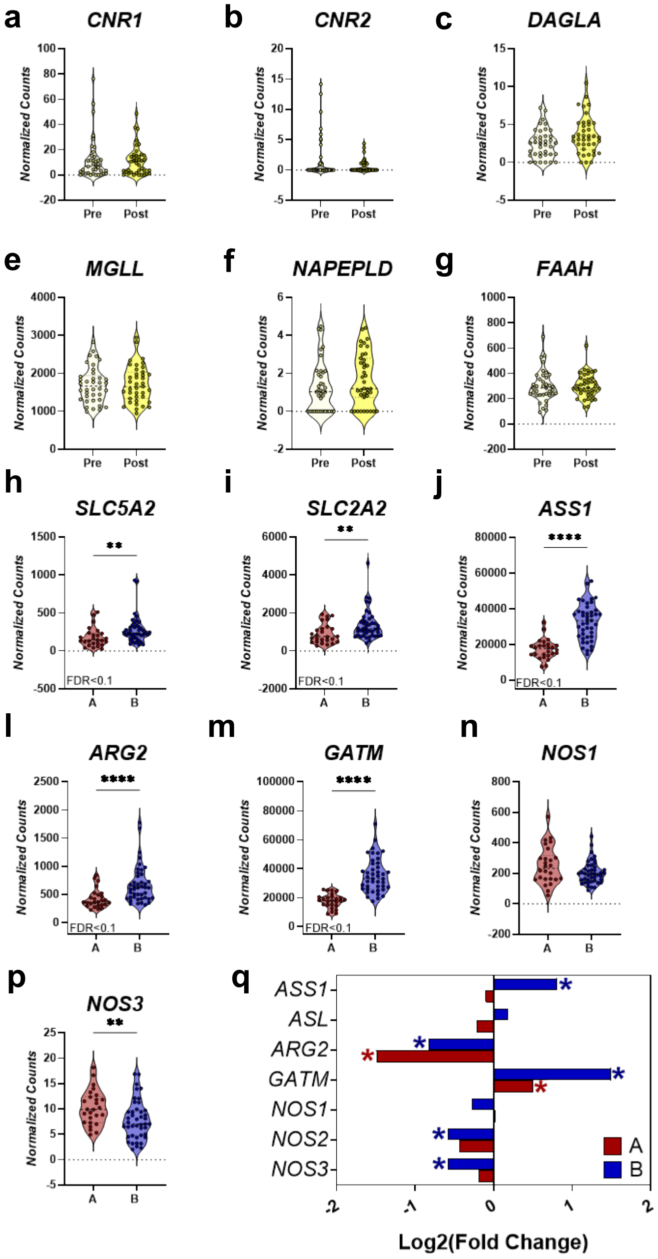


**Supplementary Figure 1. Gene expression disruptions in the kidneys of KTRs at acute and chronic disease stages.** RNA-Seq of whole kidney mRNA from 42 KTRs at multiple time points (GSE126805). **(a-g)** Gene expression immediately before (Pre) and after (Post) renal reperfusion, compared using the Mann-Whitney test (*p < 0.05, **p < 0.01, ***p < 0.001, ****p < 0.0001). Differential expression was also assessed by DESeq2 (FDR < 0.1);

the genes shown here were not differentially expressed at this acute time point. **(h-p)** At 3- and 12-month post-transplant, patients were stratified into two transcriptional communities: Community A (red), characterized by injury, inflammation, and fibrosis, and Community B (blue), showing healthy recovery, as described by *Cippa et al., 2018*[1]. Selected genes were compared between communities using the Mann-Whitney test (*p < 0.05, **p < 0.01, ***p < 0.001, ****p < 0.0001), and genes identified as differentially expressed by DESeq2 (FDR < 0.1) are indicated. **(q)** Changes in gene expression at 3 and 12 months relative to Post samples within each community; *FDR < 0.1. *Abbreviations:* KTRs, kidney transplant recipients; *CNR1/2*, cannabinoid-1/-2 receptor; *DAGLA/B*, diacylglycerol lipase A/B; *MGLL*, monoacylglycerol lipase; *NAPEPLD*, *N*-Acyl Phosphatidylethanolamine phospholipase D; *FAAH*, fatty acid amide hydrolase; *SLC5A2*, sodium glucose transporter 2 (SGLT2); *SLC2A2*, glucose transporter 2 (GLUT2); *ASS1*, arginosuccinate synthase-1; *ASL*, arginosuccinate lyase; *ARG2*, arginase-2; *GATM*, L-Arginine:glycine amidinotransferase; *NOS1/2/3*, nitric oxide synthase-1/2/3.

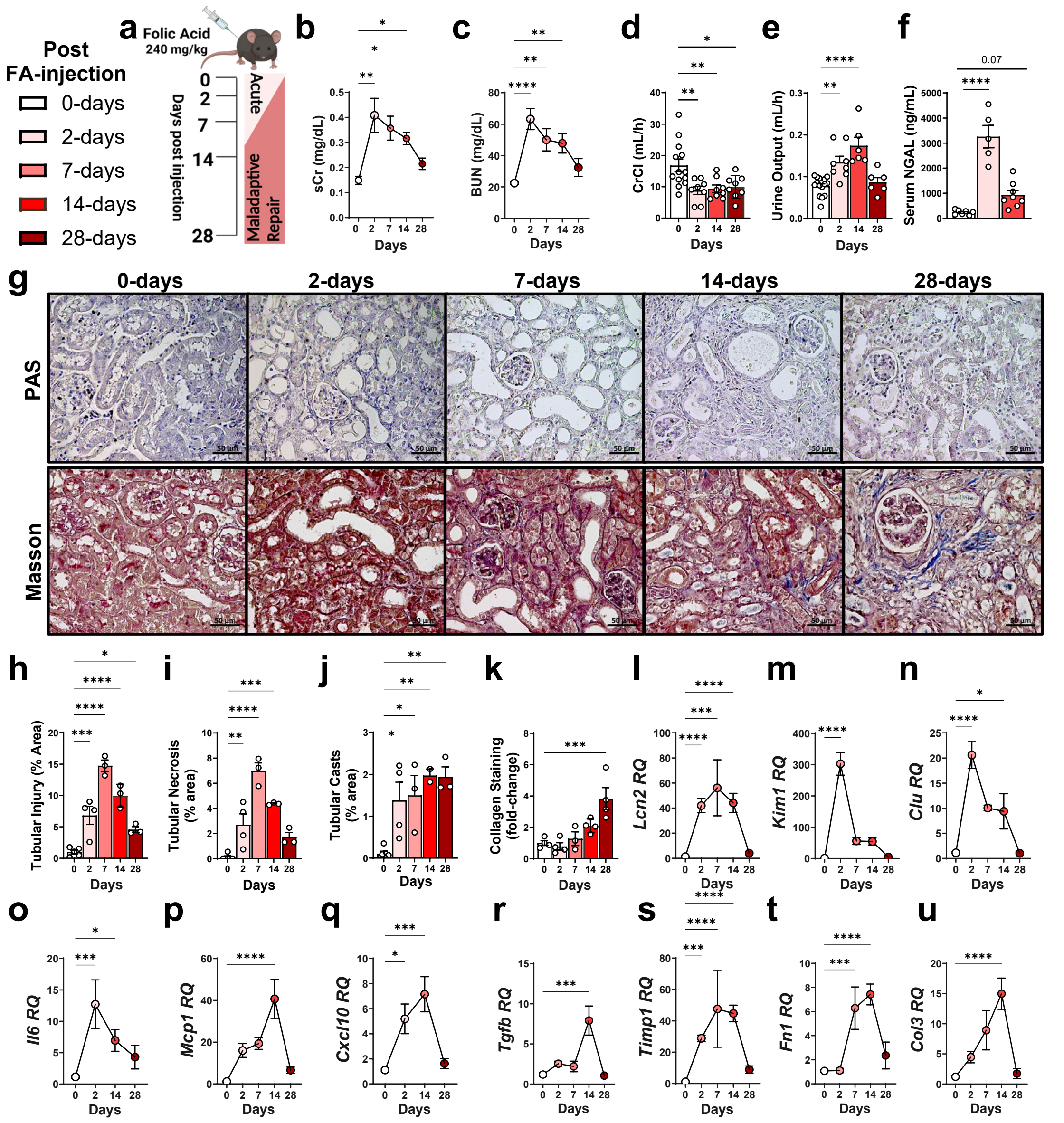


**Supplementary Figure 2. FA-induced AKI and maladaptive repair stages display distinct phenotypic signatures.** Male C57Bl/6J mice were injected intraperitoneally (i.p.) with FA (240 mg/kg) to induce AKI and evaluated at 2 days post-injection (acute injury) and at 7, 14, and 28 days (maladaptive repair stages). **(a)** Experimental timeline. **(b-d)**

Biochemical assessment of kidney function: **(b)** sCr, **(c)** BUN, **(d)** CrCl, and **(e)** rate of urinary output (n = 5-13 mice per group). **(f)** Serum NGAL levels (n = 5-8 mice per group). **(g)** Representative kidney histopathology by PAS and Masson's trichrome staining; **(h-k)** quantification of **(h)** overall tubular injury, **(i)** tubular necrosis **(j)** tubular casts formation, and **(k)** collagen (blue) staining, respectively (Scale bar: 50 µm; 40× magnification n = 3-5 mice per group). **(l-u)** Renal gene expression of markers of tubular injury, inflammation, and fibrosis. Data represent the mean ± SEM. One-way ANOVA with Dunnett's post-hoc test versus pre-FA baseline was used. *p < 0.05, **p < 0.01, ***p < 0.001, ****p < 0.0001. *Abbreviations:* FA, folic acid; AKI, acute kidney injury; sCR, serum creatinine; BUN, blood urea nitrogen; CrCl, creatinine clearance rate; NGAL, neutrophil gelatinase-associated lipocalin; PAS, Periodic Acid-Schiff; *Lcn2*, neutrophil gelatinase-associated lipocalin; *Kim1*, kidney injury marker-1; *Clu*, clusterin; *Il6*, interleukin-6; *Mcp1*, monocyte chemoattractant protein-1; *Cxcl10*, interferon gamma-induced protein 10; *Timp1*, TIMP metallopeptidase inhibitor 1; *Tgfb*, transforming growth factor beta; *Fn1*, fibronectin; *Col3*, collagen-3

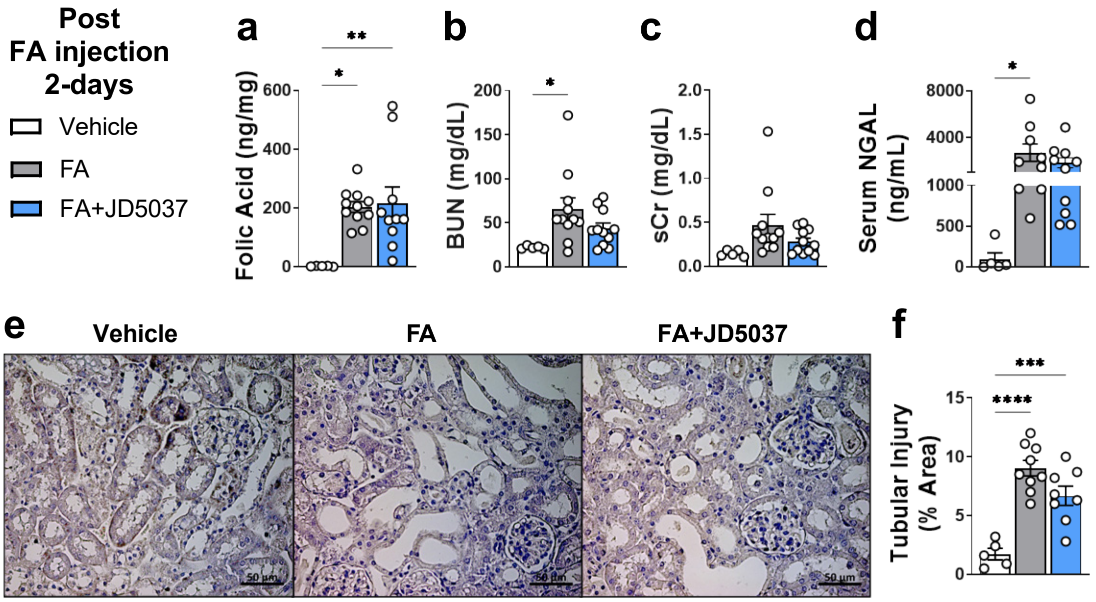


**Supplementary Figure 3. Peripheral CB1R blockade does not alter early FA-induced AKI severity.** Male C57BL/6J mice received FA (240 mg/kg, i.p.) to induce AKI or vehicle (0.3 M sodium bicarbonate), followed by 2-days of daily oral treatment with the peripherally restricted CB1R inverse agonist JD5037 (3 mg/kg) or vehicle (n = 5-11 per group). **(a)** Folic acid levels in kidney tissue. Serum levels of **(b)** BUN, **(c)** sCr, and **(d)** NGAL as indices of early kidney dysfunction and injury. **(e)** Representative kidney histopathology Periodic-Schiff (PAS) staining (Scale bar: 50μm, 40× magnification). **(f)** Quantification of relative tubular injury area (n = 5-10 per group). Data represent the mean ± SEM. One-way ANOVA with Tukey's post-hoc test was used for group comparisons. *p < 0.05, **p < 0.01, ***p < 0.001, ****p < 0.0001. *Abbreviations:* CB1R, cannabinoid-1 receptor; FA, folic acid; AKI, acute kidney injury; BUN, blood urea nitrogen; sCR, serum creatinine; NGAL, neutrophil gelatinase-associated lipocalin.

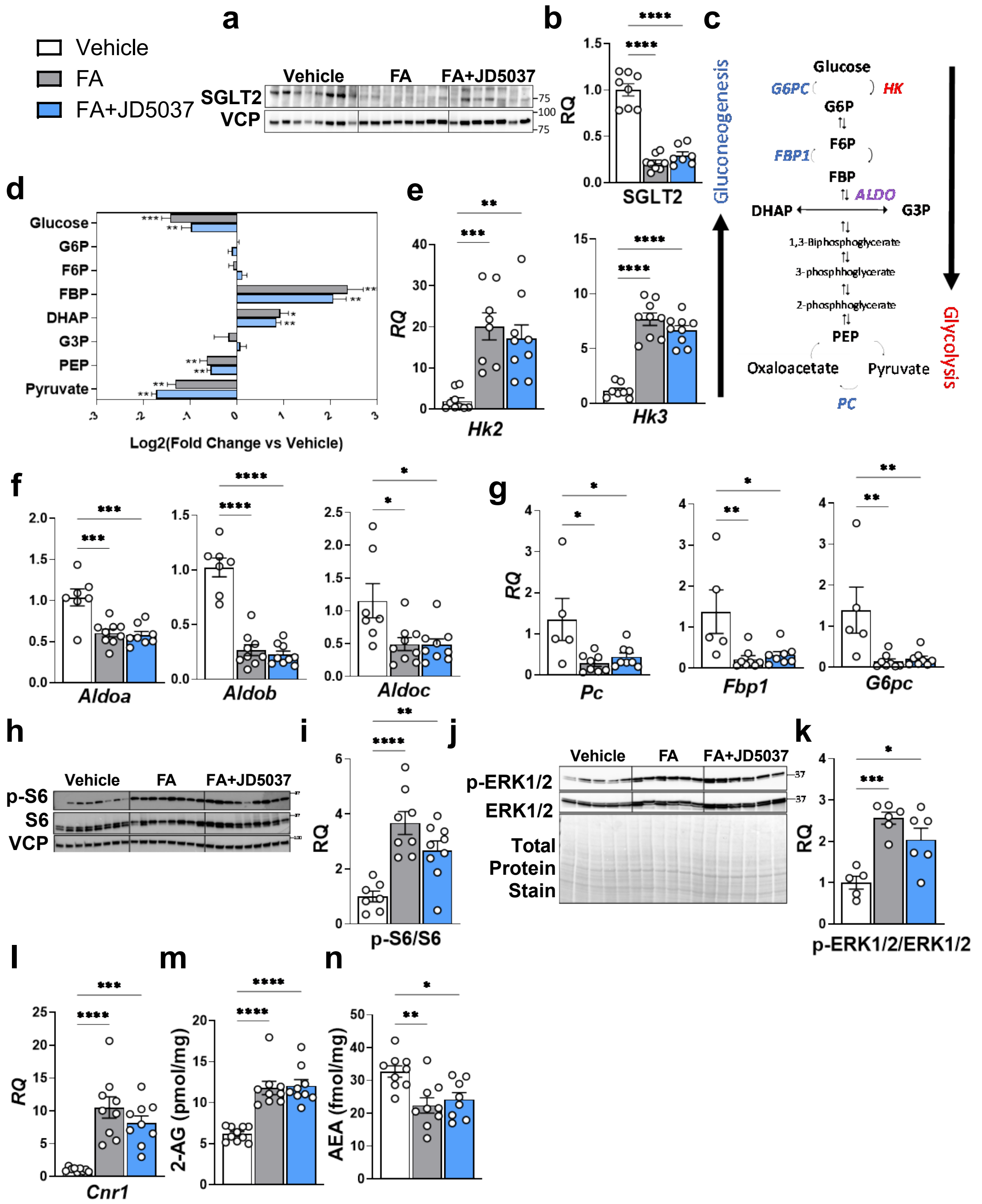


**Supplementary Figure 4. Peripheral CB1R blockade post FA-AKI does not affect renal glucose handling and signaling pathways.** Male C57BL/6J mice received daily oral treatment with the peripherally restricted CB1R inverse agonist JD5037 (3 mg/kg) or vehicle for 14 days following FA-induced AKI (240 mg/kg) or vehicle injection (0.3 M sodium bicarbonate) (n = 9-10 per group). Renal SGLT2 analysis (n = 7-8 per group): **(a)** immunoblot and **(b)** relative quantification normalized to VCP. **(c)** Schematic of glycolysis and gluconeogenesis pathways, highlighting key gluconeogenic (blue), glycolytic (red), and shared (purple) enzymes. **(d)** Fold-change of kidney metabolites in the FA-group and the FA+JD5037 groups relative to the vehicle controls (n = 6 per group). Renal transcript expression of key enzymes in **(e)** glycolysis, **(f)** shared pathways, and **(g)** gluconeogenesis (n = 5-9 per group). **(h)** Immunoblot for phosphorylated S6 and total S6 and **(i)** pS6/S6 quantification. **(j)** Immunoblot for phosphorylated ERK1/2 and total ERK1/2 and **(k)** p-ERK1/2/ERK1/2 quantification. **(l)** Renal *Cnr1* transcript expression and **(m, n)** kidney endocannabinoid levels of 2-AG and AEA. Data represent the mean ± SEM. One-way ANOVA with Tukey's post-hoc test was used for group comparisons. *$p < 0.05$, **$p < 0.01$, ***$p < 0.001$, ****$p < 0.0001$. *Abbreviations:* SGLT2, sodium glucose transporter 2; VCP, Valosin-containing protein; G6P, glucose-6-phosphate; F6P, fructose-6-phosphate; FBP, fructose-1,6-biphosphate; DHAP, dihodroxyacetone phosphate; G3P, glyceraldehyde-3-phosphate; PEP, phosphoenolpyruvate; HK2/3, hexokinase; G6PC, glucose-6-phosphatase; FBP1, Fructose biphsophatase-1; PC, Pyruvate carboxylase; ALDOa/b/c, Aldolase; p-S6/S6, phosphorylated ribosomal protein S6; p-ERK1/2/ERK1/2, phosphorylated extracellular signal-regulated kinase; *Cnr1,* Cannabinoid-1 receptor; 2-AG, 2-arachidnoylglycerol; AEA, anandamide.

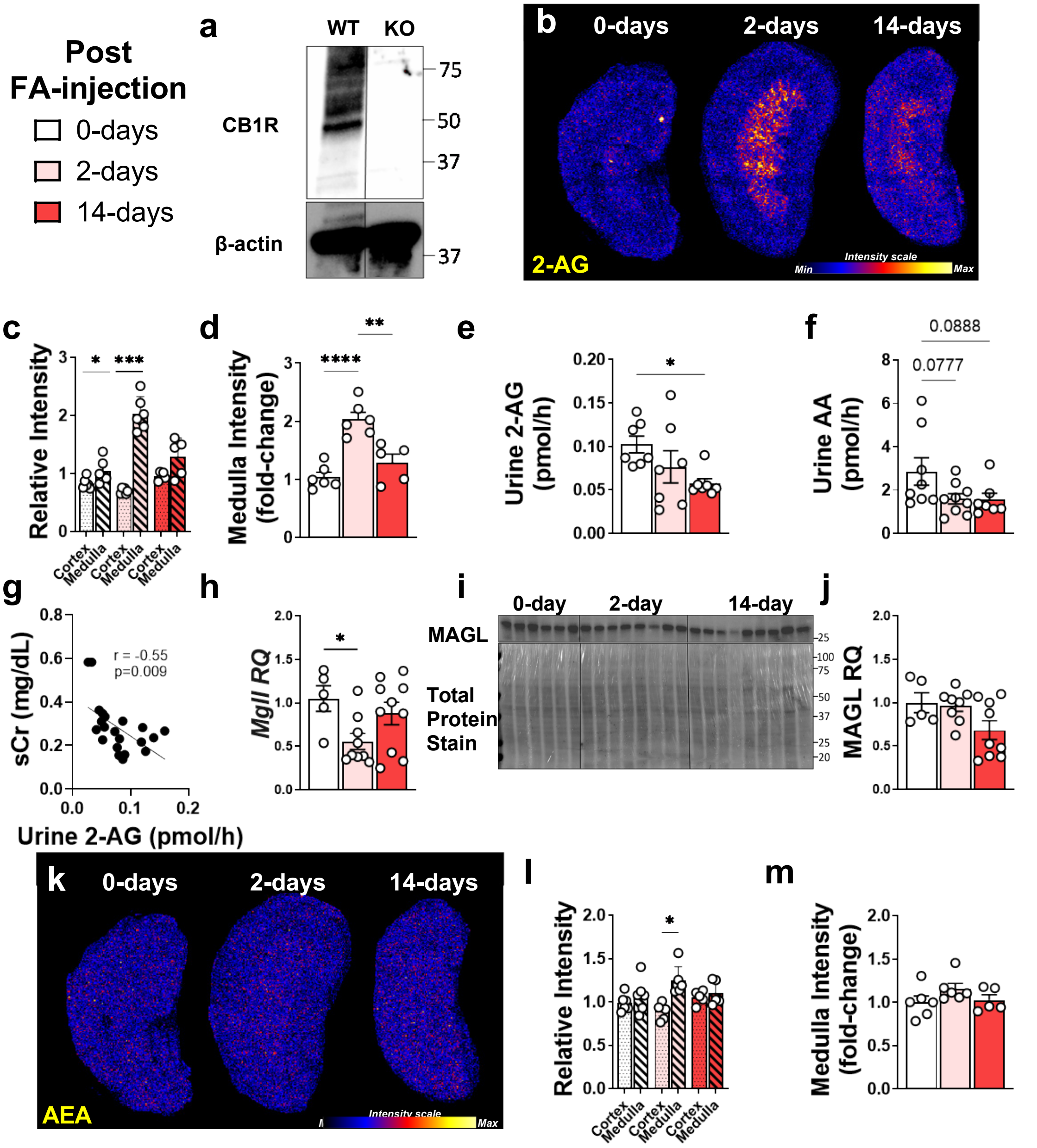


**Supplementary Figure 5. ECS changes in rodent models of AKI and maladaptive repair.** **(a)** Validation of CB1R antibody specificity using brain tissue lysates from wild-type (WT) and CB1R knockout (KO) mice. Male C57Bl/6J mice were injected with FA (240 mg/kg, i.p.) to induce AKI and evaluated at baseline (0 days), and at 2 and 14 days post-injection. **(b)** DESI-MSI imaging of 2-AG in kidney sections and **(c)** quantification of relative 2-AG intensity in renal cortex versus medulla at each time point, with **(d)** ROI highlighting the medulla (n = 5-6 per group). **(e-f)** Quantification of 2-AG and AA in 24-hour urine samples at each time point (n = 7-9 per group). **(g)** Univariate linear regression analysis with Pearson correlation coefficient between sCr and urinary 2-AG. Renal *Mgll*/MAGL analysis: **(h)** transcript expression and **(i)** Western blot, and **(j)** relative protein quantification normalized to total protein (n = 6-10 per group). **(k)** DESI-MSI imaging of AEA in kidney sections and **(l)** quantification of relative AEA intensity in cortex versus medulla at each time point, with **(m)** ROI highlighting the medulla (n = 5-6 per group). Data represent the mean ± SEM. Student's paired t-test (medulla versus cortex) or one-way ANOVA with Tukey's post-hoc test between groups was used as appropriate. *p < 0.05, **p < 0.01, ***p < 0.001, ****p < 0.0001. *Abbreviations:* CB1R, cannabinoid-1 receptor; FA, folic acid; AKI, acute kidney injury; 2-AG, 2-arachidonoylglycerol; *Mgll*/MAGL, monoacylglycerol lipase; AA, arachidonic acid; AEA, anandamide; ROI, region of interest.

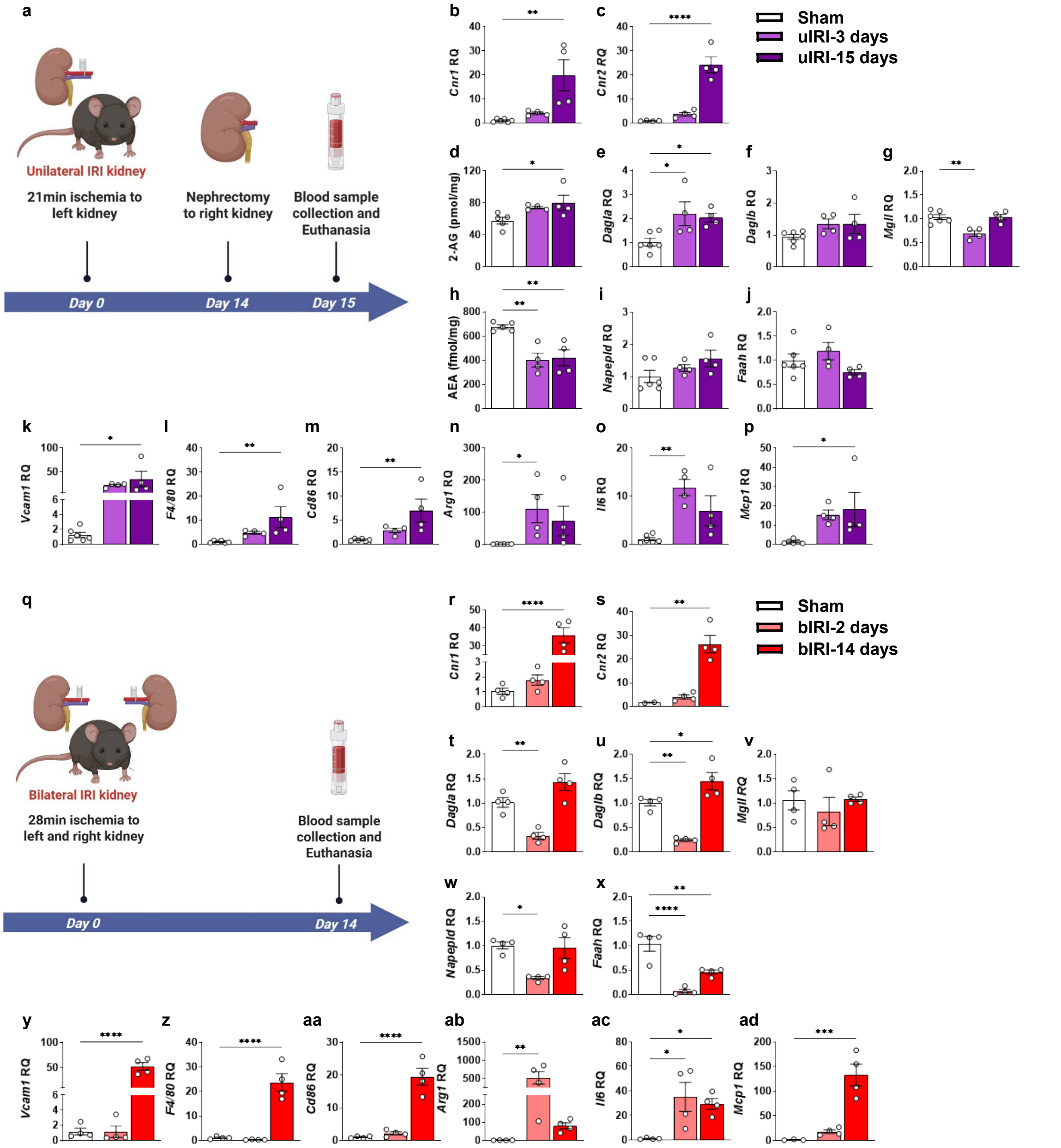


**Supplementary Figure 6. Enhanced CB1R expression and activation in murine IRI models. (a)** C57Bl/6J, 12-13 weeks old, male mice underwent unilateral IRI with delayed nephrectomy after 14 days (uIRI) and were compared to sham operation control mice (Sham: $n$ = 6, uIRI: $n$ = 4 mice per group). Whole kidney transcriptional levels of the cannabinoid receptors **(b)** *Cnr1* and **(c)** *Cnr2*. **(d)** Kidney 2-AG levels normalized to protein content. **(e, f)** The transcriptional levels of 2-AG-metabolizing enzymes (*Dagla* and

*Daglb*) and **(g)** its degrading enzyme (*Mgll*). **(h)** Kidney AEA levels normalized to protein content. **(i)** The transcriptional levels of AEA-metabolizing enzyme (*Napepld*) and **(j)** its degrading enzyme (*Faah*). **(k-p)** Renal gene expression of markers of tubular injury, inflammation, and fibrosis. **(q)** C57Bl/6J, 12-13 weeks old, male mice underwent bilateral IRI (bIRI) and were compared to sham operation control mice (Sham: *n* = 3-4, bIRI: *n* = 4 mice per group). Whole kidney transcriptional levels of the cannabinoid receptors **(r)** *Cnr1* and **(s)** *Cnr2*, **(t, u)** 2-AG metabolizing enzyme (*Dagla* and *Daglb*), and **(v)** its degrading enzyme (*Mgll*). **(w)** The transcriptional levels of AEA-metabolizing enzyme (*Napepld*) and **(x)** its degrading enzyme (*Faah*). **(y-ad)** Renal gene expression of markers of tubular injury, inflammation, and fibrosis. Data represent the mean ± SEM. One-way ANOVA with Dunnett's post-hoc test was used for group comparisons. *p < 0.05, **p < 0.01, ***p < 0.001, ****p < 0.0001. *Abbreviations*: uIRI, unilateral ischemia-reperfusion injury; bIRI, bilateral ischemia-reperfusion injury; *Cnr1*, cannabinoid 1 receptor; *Cnr2*, cannabinoid 2 receptor; *Dagla/b*, 2-AG synthesizing enzyme diacylglycerol lipase isoforms a or b; *Mgll*, 2-AG catabolic enzyme monoacylglycerol lipase; *Napepld*, AEA synthesizing enzyme *N*-acyl phosphatidylethanolamine phospholipase D; *Faah*, AEA catabolic enzyme fatty acid amide hydrolase; *Vcam1*, vascular cell adhesion molecule 1; *F4/80*, *Adgre1*- adhesion G protein-coupled receptor E1; *Cd86*, CD86 antigen; *Arg1*, arginase 1; *Il6*, interleukin-6; *Mcp1*, *Ccl2*- C-C motif chemokine ligand 2.

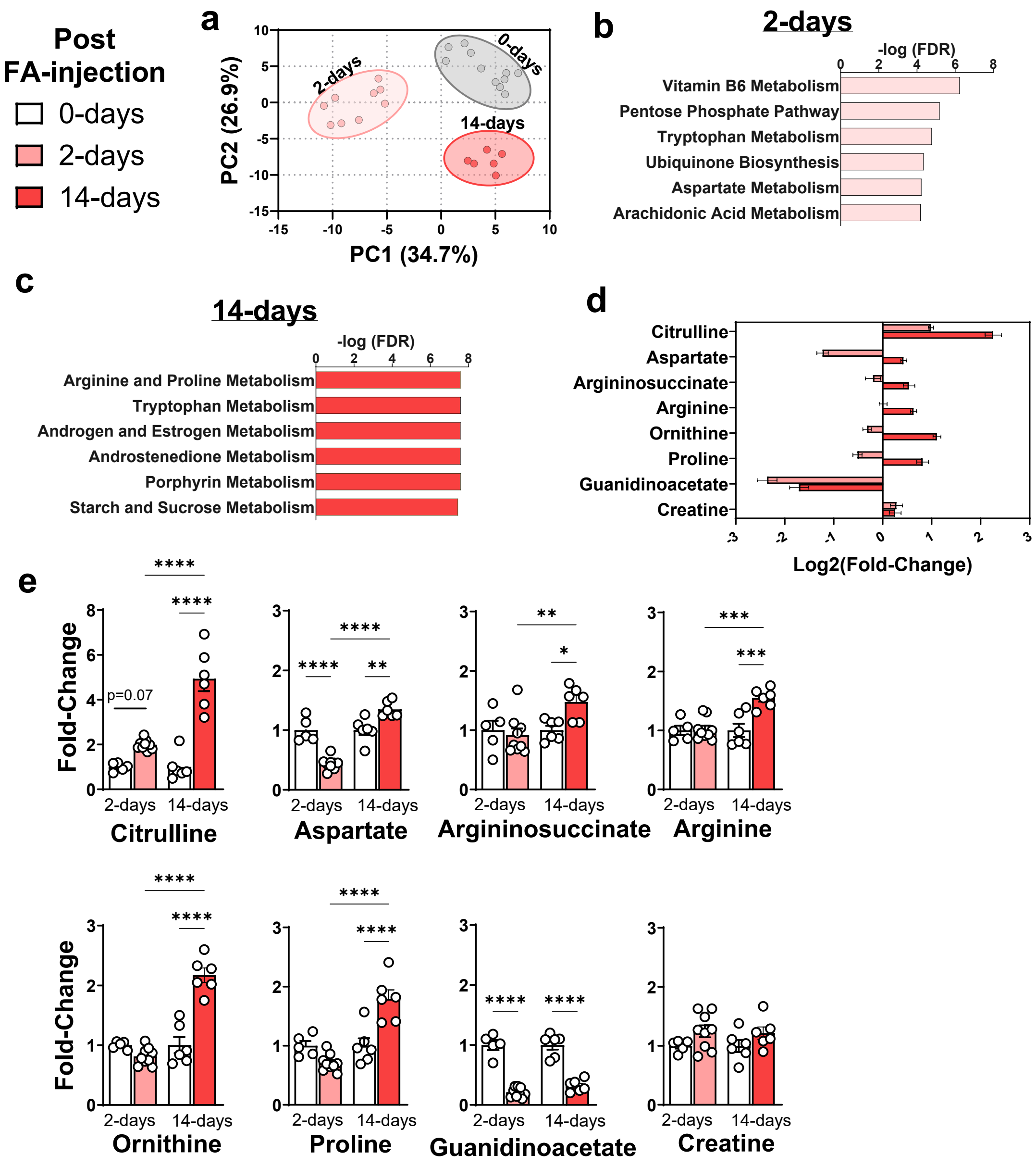


**Supplementary Figure 7. Differential changes in arginine-related metabolites in FA-induced AKI and maladaptive recovery. (a)** Kidney metabolomic profiling at 0-, 2-, and 14-day post-FA injection (n = 5-9 mice per group) with PCA illustrating separation of samples by time point. **(b, c)** Pathway enrichment analysis of differentially abundant metabolites at 2- and 14-days compared to the pre-injection (0-day) state. **(d, e)** LC-MS/MS analysis of arginine-related precursors and metabolites at 2- and 14-days following FA-induced AKI (240 mg/kg, i.p.), shown as fold-change relative to corresponding controls (n = 6 per group). Data represent the mean ± SEM. One-way ANOVA with Tukey's post-hoc test was performed. *p < 0.05, **p < 0.01, ***p < 0.001, ****p < 0.0001. *Abbreviations:* FA, folic acid; AKI, acute kidney injury; PCA, principal component analysis.

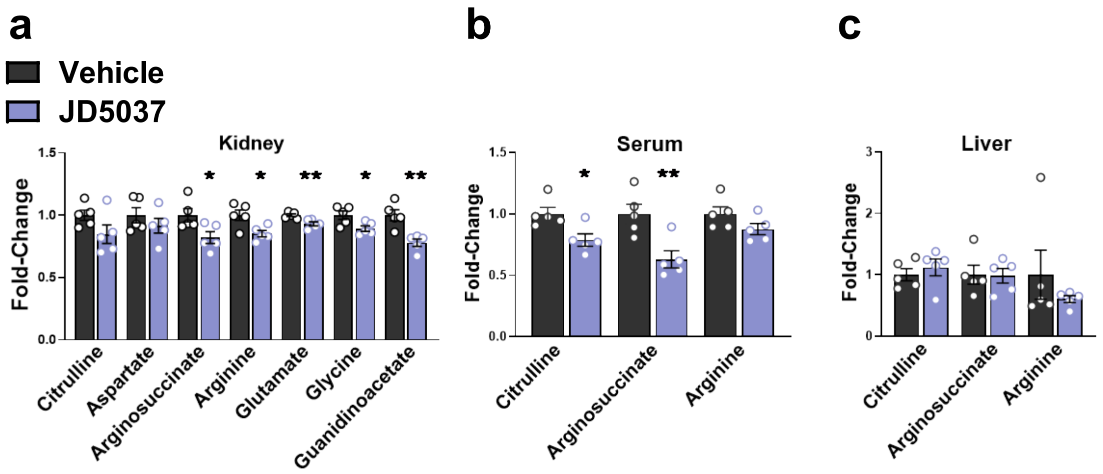


**Supplementary Figure 8. Peripheral CB1R blockade in healthy mice modulates arginine-related metabolites in serum and kidney, but not liver.** Healthy 10-12-week-old male mice were treated daily with JD5037 (3 mg/kg, i.p.) or vehicle for one week and subjected to LC-MS/MS-based profiling of arginine-related metabolites (n = 5 per group). Fold-change in metabolite levels relative to vehicle-treated controls is shown for **(a)** kidney, **(b)** serum, and **(c)** liver samples. Data represent the mean ± SEM. Student's t-test was used to compare the two groups. *p < 0.05, **p < 0.01. *Abbreviations:* CB1R, cannabinoid-1 receptor; FA, folic acid; AKI, acute kidney injury.

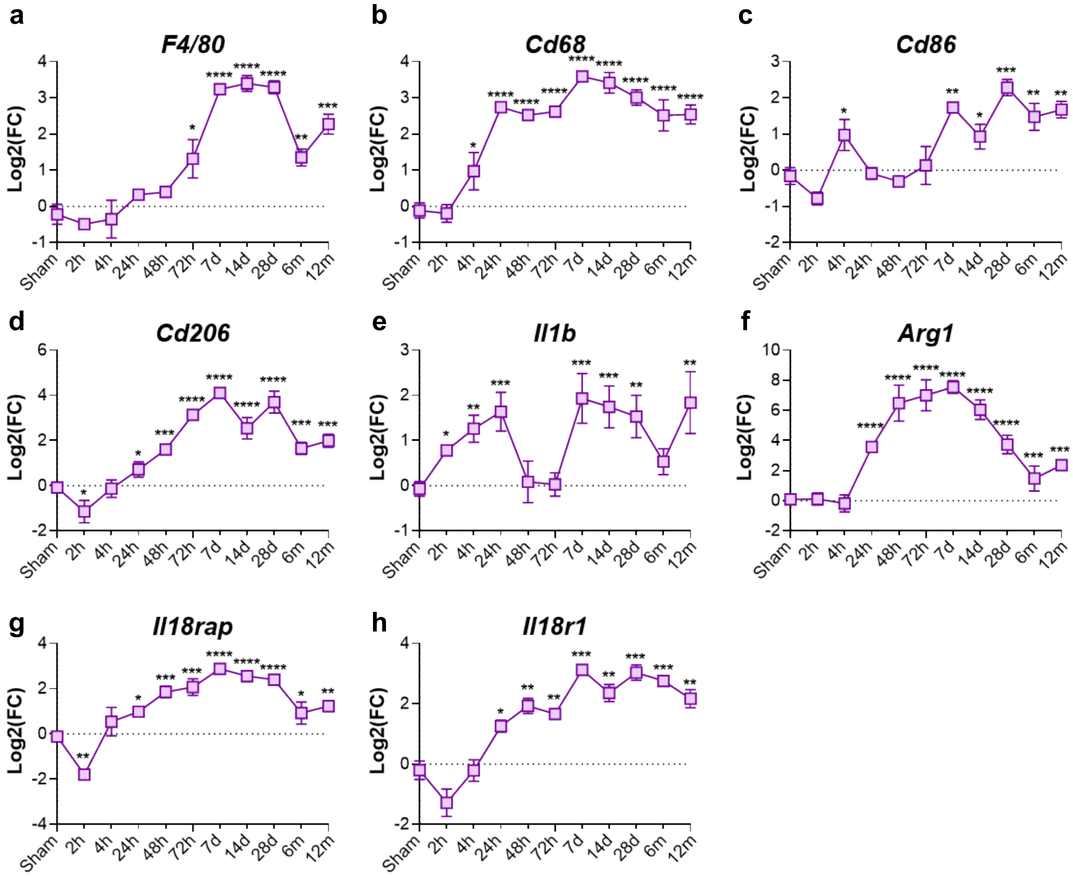


**Supplementary Figure 9. Enhanced macrophage expression and activation in post-AKI maladaptive repair.** Mouse kidneys following bilateral IRI were collected at various time points as described in Liu et al., 2017[2]. RNA-sequencing of whole mouse kidney at multiple time points after bilateral IRI (*n* = 3-4/group) compared to sham control (analyzed at 4-hour, 24-hour, 12-month; *n*=9) from NCBI GEO accession GSE98622 showing **(a-h)** macrophage-related expression and activation markers. Data represent the mean ± SEM. Comparisons by one-way ANOVA with Dunnett's post-hoc test vs sham (*p < 0.05, **p < 0.01, ***p < 0.001, ****p < 0.0001). *Abbreviations: F4/80*, *Adgre1*- adhesion G protein-coupled receptor E1; *Cd68*, CD68 antigen; *Cd86*, CD86 antigen; *Cd206*, *Mrc1*- mannose receptor, C type 1; *Il1b*, interleukin-1b; *Arg1*, arginase 1; *Il18rap*, interleukin 18 receptor accessory protein; *Il18r1*, interleukin 18 receptor 1.

**Supplementary Table 1. Endocannabinoid analysis following FA-induced AKI**

| Study 1 | | | Study 2 | | | Study 3 | | |
|---|---|---|---|---|---|---|---|---|
| | 2-AG | AEA | | 2-AG | AEA | | 2-AG | AEA |
| *Group* | *pmoles/ mg protein* | *fmoles/ mg protein* | *Group* | *pmoles/ mg tissue* | *fmoles/ mg tissue* | *Group* | *pmoles/ mg tissue* | *fmol/mg tissue* |
| 0-days | 11.0 | 71.0 | 0-days | 7.3 | 8.1 | 0-days | 5.9 | 24.5 |
| | 16.2 | 74.4 | | 8.7 | 8.9 | | 7.1 | 28.7 |
| | 22.8 | 87.5 | | 6.4 | 7.2 | | 5.2 | 28.6 |
| | 6.8 | 119.2 | | 7.8 | 7.6 | | 7.1 | 34.5 |
| | 10.0 | 69.4 | | 6.7 | 6.8 | | 5.3 | 35.7 |
| 2-days | 23.2 | 71.2 | | 7.1 | 6.2 | | 5.8 | 62.7 |
| | 11.3 | 54.3 | | 7.6 | 7.3 | | 7.2 | 31.0 |
| | 8.9 | 34.7 | 2-days | 5.9 | 5.0 | | 5.0 | 33.5 |
| | 12.3 | 48.0 | | 5.6 | 3.1 | | 7.1 | 35.9 |
| | 11.2 | 46.2 | | 4.9 | 3.5 | | 6.4 | 42.2 |
| | 7.5 | 32.6 | | 4.9 | 3.8 | 14-days | 9.7 | 20.6 |
| 7-days | 14.6 | 36.0 | | 5.5 | 5.2 | | 11.6 | 27.8 |
| | 22.2 | 40.8 | | 5.4 | 4.7 | | 11.1 | 19.1 |
| | 22.4 | 31.9 | | 6.3 | 5.0 | | 12.0 | 16.1 |
| | 11.0 | 35.7 | | 6.3 | 4.8 | | 10.2 | 36.2 |
| | 18.8 | 48.9 | | 5.9 | 4.0 | | 12.2 | 21.4 |
| | 13.8 | 39.6 | | 5.2 | 4.7 | | 10.1 | 12.4 |
| 14-days | 16.1 | 41.5 | 14-days | 7.9 | 7.6 | | 11.1 | 21.6 |
| | 19.4 | 68.6 | | 9.3 | 7.0 | | 18.0 | 26.3 |
| | 20.2 | 46.5 | | 10.0 | 3.4 | | | |
| | 25.6 | 58.9 | | 9.1 | 4.4 | | | |
| | 20.3 | 64.2 | | 9.9 | 3.4 | | | |
| | 29.1 | 40.0 | | 10.3 | 3.6 | | | |
| 28-days | 15.6 | 47.3 | | 12.9 | 8.1 | | | |
| | 11.6 | 81.0 | | 8.9 | 6.4 | | | |
| | 24.9 | 74.0 | | 9.9 | 5.3 | | | |
| | 13.7 | 62.7 | | | | | | |
| | 13.1 | 41.8 | | | | | | |
| | 28.8 | 88.2 | | | | | | |

**Supplemental Table 2. LC-MS/MS instrument settings for endocannabinoid analysis**

| Analyte | Molecular Ion $[M + H]^+$ $[M - H]^-$ for AA [m/z] | Fragment [m/z] | DP [volts] | CE [volts] | CXP [volts] |
|---|---|---|---|---|---|
| 2-AG | 379.2 | 287.1 (quantifier) | 70 | 19 | 14 |
| | | 91 (qualifier) | 70 | 67 | 10 |
| AEA | 348.2 | 287.1 (quantifier) | 26 | 13 | 16 |
| | | 62 (qualifier) | 26 | 13 | 8 |
| AA | 305.3 | 91 (quantifier) | 1 | 49 | 10 |
| | | 287.1 (qualifier) | 1 | 13 | 22 |
| $d_4$-AEA | 352.3 | 287.1 (quantifier) | 66 | 15 | 20 |
| | | 66 (qualifier) | 66 | 21 | 8 |
| d5-AG | 384.3 | 287.1 (quantifier) | 26 | 19 | 14 |
| | | 91 (qualifier) | 26 | 67 | 10 |

**Supplemental Table 3. Mouse primer sequences used for qPCR**

| Gene | Forward primer (5'-3') | Reverse primer (5'-3') |
|---|---|---|
| ***Aldoa*** | CACGAGACACTGTACCAGAAG | CACCACACCCTTATCTACCTTAAT |
| ***Aldob*** | CCAGCCTTGCTATCCAAGAA | GCACCTCTGGCTCAACAATA |
| ***Aldoc*** | GGGTCATCTTCTTCCATGAGAC | ACCTTGATGCCTACGAGAATG |
| ***Clu*** | AGAGCTCACCCTTCTACTTCTG | TCCACCTTCTCTTAAGAAATCAAC |
| ***Cnr1*** | CCGCAAAGATAGTCCCAATG | AACCCCACCCAGTTTGAAC |
| ***Cnr2*** | CTGCAGCTCTTGGGACCTAC | TGTCCCAGAAGACTGGGTGT |
| ***Col3*** | GCCCACAGCCTTCTACAC | CCAGGGTCACCATTTCTC |
| ***Cxcl10*** | GGATGGCTGTCCTAGCTCTG | TGAGCTAGGGAGGACAAGGA |
| ***Dagla*** | AGAATGTCACCCTCGGAATG | GGTCTTCCTGTAGCTGTGGGCC |
| ***Faah*** | GTATCGCCAGTCCGTCATTG | GCCTATACCCTTTTTCATGCCC |
| ***Fbp1*** | GCATCGCACAGCTCTATGGT | TTGGATGAGCCATCAAGGGG |
| ***Fn1*** | ATGTGGACCCCTCCTGATAGT | GCCCAGTGATTTCAGCAAAGG |
| ***G6pc*** | TTTCCCCACCAGGTCGTGGCT | CCCATTCTGGCCGCTCACACC |
| ***Gapdh*** | AGGTCGGTGTGAACGGATTTG | TGTAGACCATGTAGTTGAGGTCA |
| ***Hk2*** | GGTACAGAGAAAGGAGACTTC | TCTTGTTATGCATCTCTACGC |
| ***Hk3*** | CACTTAACCAATCTCGGAGT | AGGCTATCACTTTCGATCTC |
| ***Il18*** | GACTCTTGCGTCAACTTCAAGG | CAGGCTGTCTTTTGTCAACGA |
| ***Il6*** | GACAACCACGGCCTTCCCTA | GCCTCCGACTTGTGAAGTGGT |
| ***Kim1*** | TGTCGAGTGGAGATTCCTGGATGGT | GGTCTTCCTGTAGCTGTGGGCC |
| ***Lcn2*** | AAACAGAAGGCAGCTTTACGA | TCTGATCCAGTAGCGACAGC |
| ***Mcp1*** | GCATTAGCTTCAGATTTA | TTAAAAACCTGGATCGGAACCAA |
| ***Mgll*** | ACCATGCTGTGATGCTCTCTG | CAAACGCCTCGGGGATAACC |
| ***Napepld*** | ACGTCCTCCTCTAGTCTGTAATC | AGCGCCAAGCTATCAGTATCC |
| ***Pc*** | GATGACCTCACAGCCAAGCA | GGGTACCTCTGTGTCCAAAGGA |
| ***Rps16*** | AGGAGCGATTTGCTGGTGTGG | GCTACCAGGGCCTTTGAGATG |
| ***Tgfb*** | GCGGACTACTATGCTAAAGAGG | GTAGAGTTCCACATGTTGCTCC |
| ***Timp1*** | TGTGGGAAATGCCGCAGATA | TTCACTGCGGTTCTGGGACT |

**Supplemental Table 4. Human primer sequences used for qPCR**

| Gene | Forward primer (5'-3') | Reverse primer (5'-3') |
|---|---|---|
| ***ICAM1*** | GGCCGGCCAGCTTATACAC | TAGACACTTGAGCTCGGGCA |
| ***MCP1*** | ATGCAATCAATGCCCCAGTC | TGCAGATTCTTGGGTTGTGG |
| ***TNFa*** | CCCTGGTATGAGCCCATCTATC | AAAGTAGACCTGCCCAGACTCG |
| ***VCAM1*** | CAGTAAGGCAGGCTGTAAAAGA | TGGAGCTGGTAGACCCTCG |
| ***ASS1*** | TCTACAACCGGTTCAAGGGC | TCCAGGATTCCAGCCTCGTA |
| ***ARG2*** | TCAGTGCTGCGGATCATGT | CACTCCTTTTCTTTTCTGCCCTT |
| ***CXCL10*** | TGAAATTATTCCTGCAAGCCAA | CAGACATCTCTTCTCACCCTTCTTT |
| ***IL6*** | TAGCCGCCCCACAGACAG | GGCTGGCATTTGTGGTTGGG |
| ***CNR1*** | GGGGCCTGTGAATGGATATGT | GTGTTCCACCGCAAAGATAGC |
| ***DAGLa*** | CCATCTTCCTCTTTCTCCT | CTCGTGCGGGTTATAGAC |
| ***DAGLb*** | TCAGGTGCTACGCCTTCTC | TCACACTGAGCCTGGGAATC |
| ***MGLL*** | GGAAACAGGACCTGAAGACC | ACTGTCCGTCTGCATTGAC |
| ***RPLP*** | CTTCCTTAAGATCATCCAACTA | ACATGCGGATCTGCTGCA |

**Supplementary Table 5. Antibodies used in the study**

| Target | Antibody (Product code, Source) | Dilution |
|---|---|---|
| **CB1R** | 10006590, Cayman | 1:1000 |
| **DAGLα** | ab81984, Abcam | 1:500 |
| **MAGL** | ab228598, Abcam | 1:1000 |
| **FAAH** | ab54615, Abcam | 1:1000 |
| **NGAL** | ab70287, Abcam | 1:500 |
| **SGLT2** | ab385626, Abcam | 1:1000 |
| **GLUT2** | AG022, Alomone labs | 1:500 |
| **ASS1** | ab170952, Abcam | 1:1000 |
| **ARG2** | ab228700, Abcam | 1:1000 |
| **GATM** | ab228937, Abcam | 1:1000 |
| **IKK alpha** | 11930, Cell signaling | 1:1000 |
| **IKK beta** | 8943, Cell signaling | 1:1000 |
| **Phospho-IKK alpha/beta** | 2697, Cell signaling | 1:500 |
| **Phospho-NF-kappaB p65** | 3033, Cell signaling | 1:500 |
| **IkappaB alpha** | 4814, Cell signaling | 1:1000 |
| **Phospho-IkappaB alpha** | 2859, Cell signaling | 1:500 |
| **NF-kappaB p65** | 8242, Cell signaling | 1:1000 |
| **VCP** | ab155146, abcam | 1:1000 |
| **Anti-rabbit** | ab97085, abcam | 1:1000 |
| **Anti-rat** | ab97057, abcam | 1:5000 |
| **Anti-goat** | ab97120, abcam | 1:2500 |
